# Drug-induced cis-regulatory elements in human hepatocytes affect molecular phenotypes associated with adverse reactions

**DOI:** 10.1101/2024.07.24.604883

**Authors:** Saki Gotoh-Saito, Ryoko Wada, Hideya Kawaji

## Abstract

**Background:** Genomic variations contribute to the phenotypic diversity of individuals. A number of polymorphisms in protein-coding regions that alter drug efficacy or lead to adverse reactions have been characterized; however, noncoding regions that affect drug responses are largely overlooked, except for a limited number of well-studied enhancers.

**Results:** We conducted a quantitative assessment of *cis*-regulatory elements (CREs) based on transcription initiation profiling of mRNAs and noncoding RNAs, including enhancer RNAs, by using CAGE (Cap Analysis of Gene Expression). Candidate CREs identified in a hepatocellular carcinoma HepG2 cell line with stable expression of drug-responsive transcription factor pregnane X receptor (PXR) were further narrowed down by integrating data of PXR-binding sites in human primary hepatocytes and genome-wide association studies. We found more than 100-fold enrichments of the candidates to genetically associated loci with circulating levels of bilirubin and vitamin D, which implicated a link to adverse reactions of PXR ligands. We uncovered novel enhancers of *UGT1A1* and *TSKU* through CRISPR/Cas9 knockout experiments. We identified alleles altering regulatory activities of *UGT1A1* and *CYP24A1*enhancers by using luciferase reporter assay. Furthermore, our siRNA experiments revealed an unexpected impact of TSKU on the expression of vitamin D-metabolizing enzymes.

**Conclusions:** Our transcriptome-based assessment of CREs expanded the list of drug-inducible and PXR-mediated enhancers and super-enhancers. We identified regulatory alleles that alter drug-induced gene expressions, and discovered a novel molecular cascade associated with an adverse reaction. Our results contribute a precise understanding of the noncoding elements of the human genome underlying drug responses.

## Background

Variations in the human genome contribute to the phenotypic differences among individuals. Understanding their impact on drug efficacy and adverse drug reactions (ADRs) [1] is of importance in drug development and administration. Extensive studies in the field of pharmacogenomics have been conducted over decades, leading to the discovery of causal variants that modulate drug responses. For instance, drug-metabolizing enzymes encoded by *cytochrome P450 family 2 subfamily C member 9* (*CYP2C9*) and *member19* (*CYP2C19*) harbor missense polymorphisms (CYP2C9*2, CYP2C9*3, CYP2C19*2, and CYP2C19*3) leading to reduced enzymatic activity [2–4]. Most of the genetic variants with established causal relationships to altered drug responses are located in the protein-coding regions, particularly in genes for which the contributions to drug-associated phenotypes are well studied.

Meanwhile, over 90% of the trait-associated genetic variations identified in genome-wide association studies (GWAS) are located in noncoding regions [5] [6]. These variants are shown to be significantly enriched in *cis*-regulatory elements (CREs), such as promoters and enhancers, which control gene expression either in the proximity of the target genes or from distant locations [5, 7–10]. A well-documented noncoding variant associated with a drug response is the polymorphism at the promoter region of *uridine diphosphate-glucuronosyltransferase 1A* (*UGT1A1*), known as UGT1A1*28. The allele causes a decrease of *UGT1A1* expression level [11], consequently leading to an elevated risk of neutropenia in treatment of the anticancer drug irinotecan [12]. Assessment of this polymorphism is now recommended for clinical treatment of patients [13]. This example underscores the importance of comprehending CREs in the noncoding regions of the genome.

The nuclear receptor pregnane X receptor (PXR), or NR1I2, is a drug-activated transcription factor primarily expressed in the liver and the intestine. PXR has a flexible ligand-binding pocket that allows it to be activated by a wide range of prescription drugs, including rifampicin (an antimicrobial drug), dexamethasone (an anti-inflammatory corticosteroid), phenobarbital (a sedative and anti-seizure drug for epilepsy), and tamoxifen (a selective estrogen receptor modulator to prevent breast cancer). Vitamin D deficiency is a well-documented adverse effect of PXR activators, such as phenytoin and phenobarbital [14, 15]. Upon activation by the drugs, PXR forms a heterodimer with retinoid X receptor α (or NR2B1) and binds to a specific set of CREs throughout the genome, leading to the transcription of genes that encode drug-metabolizing enzymes and drug transporters [16–18]. The well-characterized targets of PXR are xenobiotic response enhancer module (XREM) for *cytochrome P450 family 3 subfamily A member 4 (CYP3A4)* [19] and phenobarbital-responsive enhancer module (PBREM) for *UGT1A1* [20]. A minor allele in PBREM (UGT1A1*60 of rs4124874) decreases transcriptional activity of *UGT1A1* [21].

Smith *et al*. [22] conducted a genome-wide survey of drug-inducible and PXR-binding CREs in human primary hepatocytes. The survey employed epigenetic profiling, in particular ChIP-seq assays targeting PXR, p300, and histone modifications H3K4me1 and H3K27ac. Assessment of conditional occupancy of PXR and chromatin signatures led to approximately 300 potential candidates of drug induced enhancers. Of the two well-characterized enhancers, only XREM was identified. Reporter assays confirmed regulatory activities in 15 out of the 42 selected regions. This pioneering study highlights the role of enhancers in drug responses, with underscoring the need for further studies to understand enhancers altering drug responses, given the limitation of the approach based on conditional analysis of epigenetic profiles in sensitivity and specificity.

Transcriptome profiles can also be used for CRE identification, which is made possible through the detection of noncoding RNAs transcribed from active enhancers [23], called enhancer RNAs (eRNAs). Profiling of transcription start sites can be effectively used for enhancer identification [24]. This approach offers unique advantages in the assessment of regulatory activities, because eRNA level can be used as a proxy for CRE activity [25, 26] and because specificity of enhancer identification is higher in this approach than in epigenetic profiling [27]. A genome-wide atlas of promoters and enhancers in a wide range of primary cells, tissues, and cell lines has been developed in the FANTOM5 project [8, 27, 28] by using cap analysis of gene expression (CAGE), a quantitative method to monitor capped 5′ ends of RNAs [29, 30]. However, drug responsiveness in human hepatocytes was not covered.

In the present study, we aimed to delineate drug-inducible CREs mediated by PXR for a better understanding of drug responses and ADRs. We conducted a genome-wide assessment of CREs in a HepG2 cell line with stable expression of human PXR (ShP51 cells) [31] by using CAGE. By integrating data on PXR-binding sites found in human primary hepatocytes [22] and GWAS results, we narrowed down the candidate regions to the ones directly targeted by PXR and associated with phenotypes. We assessed the impact of novel enhancers on drug-responding phenotypes by using CRISPR/Cas9 knockout, luciferase reporter assay, and siRNA knockdown experiments.

## Results

### Genome-wide survey of drug-inducible CREs through quantitative profiling of transcription start sites

The human hepatocellular carcinoma cell line HepG2 is a widely used model system, but it is not necessarily suitable for studying drug responses as it is, owing to the absence of PXR expression. As an *in vitro* model of human hepatocytes, we chose ShP51 cells, which were generated from HepG2 cells to stably express human PXR [31]. We confirmed upregulation of both *CYP3A4* mRNA and XREM eRNA upon treatment with rifampicin in ShP51 cells (Fig. 1A). This demonstrated that the model system recapitulates the response to PXR activation, and that activation of both promoters and enhancers can be assessed by RNA quantification.

**Fig. 1.**
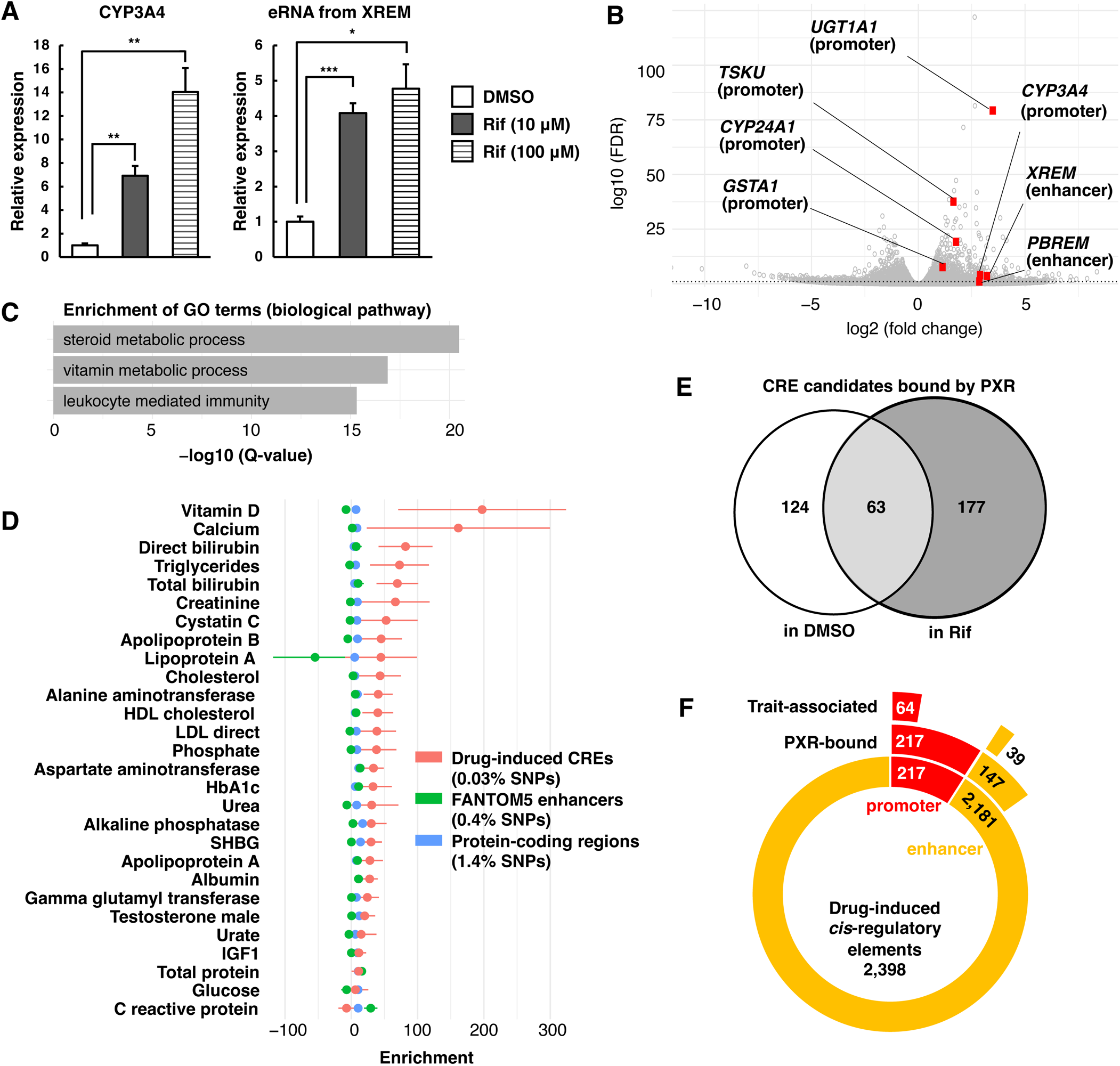
Identification of drug-inducible, PXR-binding, and trait-associated enhancer candidates in human hepatocytes. **A** qRT-PCR analysis of *CYP3A4* mRNA and xenobiotic response enhancer module (XREM) enhancer RNA (eRNA) in rifampicin (Rif)-treated ShP51 cells. The expression levels are normalized to *GAPDH* mRNA. The experiment was performed in triplicate for each condition and repeated three times with similar results. Representative result was shown. The error bars indicate standard deviation, and unpaired Welch’s *t*-test was used to calculate the *P* value. \**P* < 0.05, \*\**P* < 0.01, and \*\*\**P* < 0.001. **B** Volcano plot of the activity of the identified *cis*-regulatory elements (CREs). The dotted line indicates false discovery rate (FDR) = 0.1. **C** The top three enriched GO terms associated with genes near the induced CREs, computed by Genomic Regions Enrichment of Annotations Tool. **D** Heritability enrichment of the drug-induced CREs, FANTOM5 enhancers, and protein-coding regions to summary statistics from genome-wide association studies (GWAS) on biochemical markers measured in serum, computed by the stratified linkage-disequilibrium score regression analysis. Error bars indicate the standard error. **E** Venn diagram showing the number of drug-induced CREs that overlap with pregnane X receptor (PXR)–binding sites in human primary hepatocytes, based on previously published ChIP-seq data [22]. **F** The number of drug-induced CREs and their overlap with PXR-binding sites and with GWAS SNPs.

We next performed a genome-wide survey of the response of ShP51 cells to rifampicin treatment by using CAGE. We identified 90,871 CRE candidates and found that 2,398 (2.6%) of them were induced by rifampicin treatment with statistical significance at a false discovery rate (FDR) < 0.1. We confirmed that the identified CREs included promoters of *CYP3A4*, *UGT1A1*, and *glutathione S-transferase α 1 (GSTA1)*, as well as the well-studied enhancers, XREM and PBREM (Fig. 1B and Fig. S1). We found that 217 (9%) regions were located at promoters of known genes, and the remaining 91% were located in distal regions (Fig. 1F)

We subsequently examined the impact of the drug-induced CRE candidates by assessing the functional categories (GO terms) of their nearby genes. The analysis revealed an enrichment of relevant biological processes (Fig. 1C), such as steroid metabolic processes. The result aligns with the known role of PXR in activating *CYP3A4*, which encodes a major enzyme responsible for steroid hormone hydroxylation [32]. The enrichment in vitamin metabolic process is consistent with the known ADR of PXR activating drugs, vitamin D deficiency [14, 15]. Additionally, the enrichment of leukocyte mediated immunity correlated with the fact that ligand-activated PXR exerts immunosuppressive and anti*-*inflammatory effects by antagonizing nuclear factor κB [33] [34].

We further assessed the genetic correlation of the candidate CREs with biochemical marker–associated loci identified in UK BioBank’s GWAS studies [9, 35]. Using the stratified linkage-disequilibrium score regression analysis (S-LDSC) [36] with the summary statistics, we found that the variants associated with the levels of vitamin D and bilirubin in the serum are substantially enriched in the identified CRE candidates (Fig. 1D). The enrichment in the bilirubin level–associated loci is concordant with a PXR downstream pathway, as metabolic clearance of bilirubin is driven by *UGT1A1* expression mediated by PXR binding to both the promoter and PBREM [20]. The greatest enrichment was found in the association with vitamin D and calcium levels, which was consistent with the GO enrichment analysis of nearby genes mentioned above. The high level of enrichment, such as more than 100-fold, indicated the relevance of the identified CRE candidates as regulatory elements underlying the phenotypes.

### Drug-inducible CRE candidates bound by PXR in human primary hepatocytes

We further narrowed the CRE candidates down to the ones directly targeted by PXR in human primary hepatocytes. We re-processed the data produced by a genome-wide mapping of PXR-binding sites with ChIP-seq [22]. Of the 2,398 rifampicin-induced CRE candidates, we found the overlap with PXR-binding sites (Fig. 1E) to be 364 CREs, which included 217 promoters and 147 enhancers (Fig. 1F). We refer to these sites as drug-inducible and <u>P</u>XR-binding <u>p</u>romoters (DPP; hereafter numbered as DPP1–217) and <u>d</u>rug-inducible and <u>P</u>XR-binding <u>e</u>nhancers (DPE; hereafter numbered as DPE1–147) (Table S1).

We examined the index SNPs (single nucleotide polymorphisms) in the GWAS catalog database [6] and found them or SNPs correlated with them (linkage disequilibrium as *R*^2^ > 0.8) within 103 of the 364 regions, including 64 promoters and 39 enhancers (Fig. 1F and Table S2). Three of the enhancer candidates (DPE15–17) overlapped SNPs associated with bilirubin levels (Table 1). DPE16 corresponds to PBREM, and DPE15 and DPE17 are novel enhancer candidates located upstream and at the intron of *UGT1A1*, respectively (Fig. 2A). Six of the enhancer candidates overlapped SNPs associated with vitamin D levels, including ones located downstream of *CYP24A1* (DPE127–129, Fig. 3A) and others located upstream of *TSKU* (DPE70–72, Fig. 4A). Notably, the enhancer candidates close to *CYP24A1* and *TSKU* were located within super-enhancers (Fig. 3A and Fig. 4A).

**Table 1.**
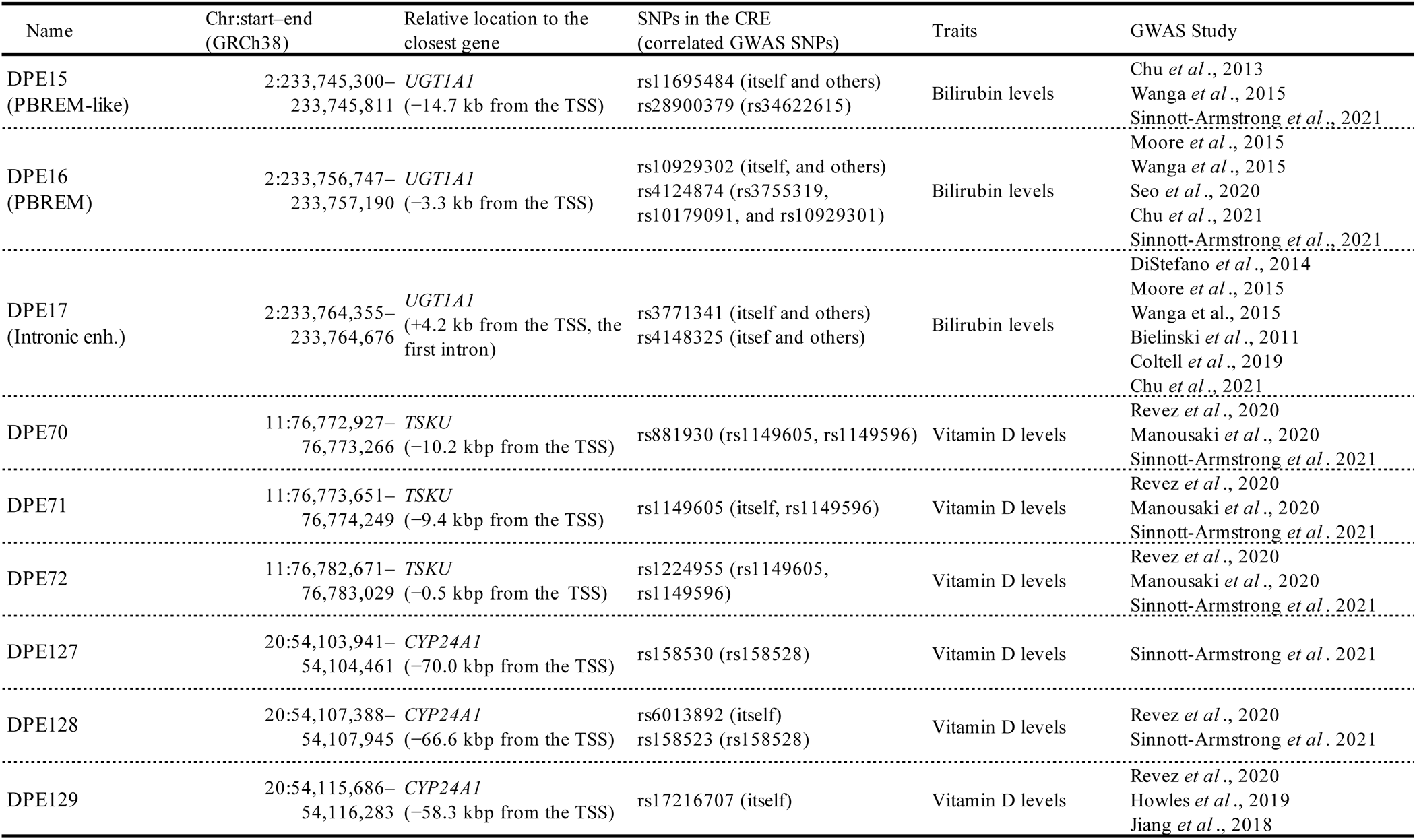
CRE candidates with SNPs associated with bilirubin and vitamin D levels.

**Fig. 2.**
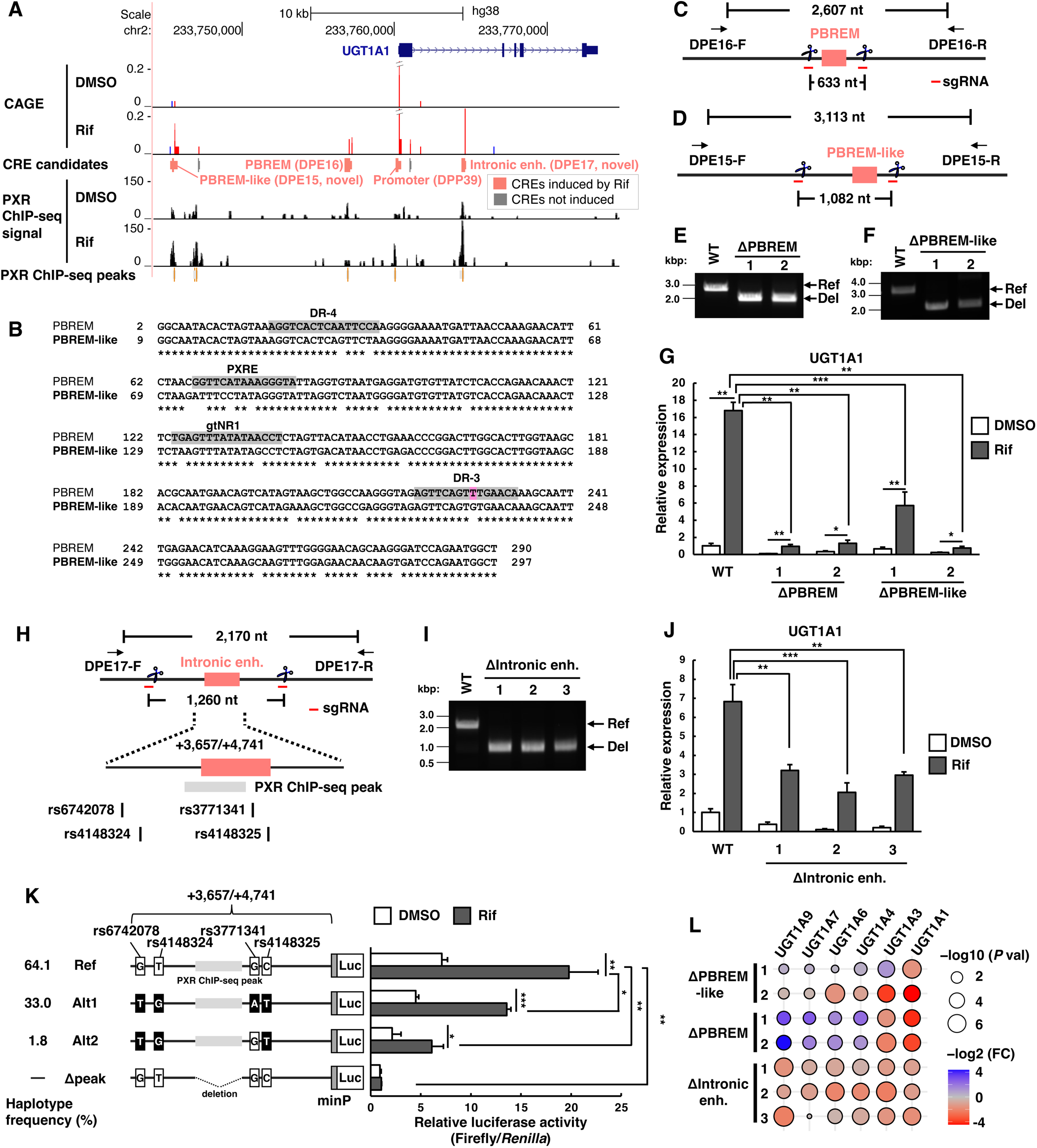
Identification of drug-inducible *UGT1A1* enhancers. **A** A view of the *UGT1A1* locus in UCSC Genome Browser. The tracks for cap analysis of gene expression (CAGE) indicate the frequencies of monitored transcription start sites at single base-pair (bp) resolution in transcripts per million, where red and blue indicate the orientation of transcription (red: forward, blue: reverse). ChIP-seq signals indicate normalized read coverage per 50 bp bin in reads per kilobase per million mapped reads. **B** Alignment of sequences for phenobarbital-responsive enhancer module (PBREM) and PBREM-like. Asterisks (*) indicate nucleotide matches. Previously reported pregnane X receptor (PXR) responsive elements in PBREM [74] are highlighted in gray, and a mismatch SNP (rs4124874, UGT1A1*60 T>G), associated with decreased activity of *UGT1A1* promoter, is indicated in pink. **C, D** Schematic view of CRISPR/Cas9-based deletion of PBREM (<u>d</u>rug-inducible and <u>P</u>XR-binding <u>e</u>nhancer (DPE)16) and PBREM-like (DPE15) in ShP51 cells with PCR primers used for amplification of genomic DNA. nt, nucleotides. **E, F** Gel images show the PCR products from genomic DNA extracted from the deletion mutants ΔPBREM 1 and 2 (**E**) and ΔPBREM-like 1 and 2 (**F**). **G** qRT-PCR analysis of *UGT1A1* mRNA normalized to *GAPDH* mRNA in rifampicin (Rif)-treated ΔPBREM and ΔPBREM-like cells. **H** Schematic view of CRISPR/Cas9-based deletion of the intronic enhancer (DPE17) in ShP51 cells. **I** The gel image shows PCR products from genomic DNA extracted from DPE17 deletion mutants (ΔIntronic enh. 1–3). **J** qRT-PCR analysis of *UGT1A1* mRNA normalized to *GAPDH* mRNA in Rif-treated ΔIntronic enh. 1–3 cells. **K** Luciferase activity of the +3,657/+4,741 region (Intronic enh.) from *UGT1A1* transcription start site with the minimal promoter (minP) in Rif-treated ShP51 cells. Three haplotypes (Ref, Alt1, and Alt2) and the same +3,657/+4,741 region lacking the PXR ChIP-seq peak (Δpeak) were assessed. Results are expressed as fold change compared with an empty vector control. Firefly luciferase activity was normalized to *Renilla* luciferase activity. **L** Effects of enhancer deletions on *UGT1A* isoform expression, where circle colors and sizes indicate fold change (FC; log^2^) and *P* values (−log^10^), respectively. Individual results are shown in Fig. S3B. All experiments were performed in triplicate or quadruplicate for each condition and repeated at least three times with similar results. Representative data were shown. The error bars indicate standard deviation, and unpaired Welch’s *t*-test was used to calculate the *P* value. \**P* < 0.05, \*\**P* < 0.01, and \*\*\**P* < 0.001.

**Fig. 3.**
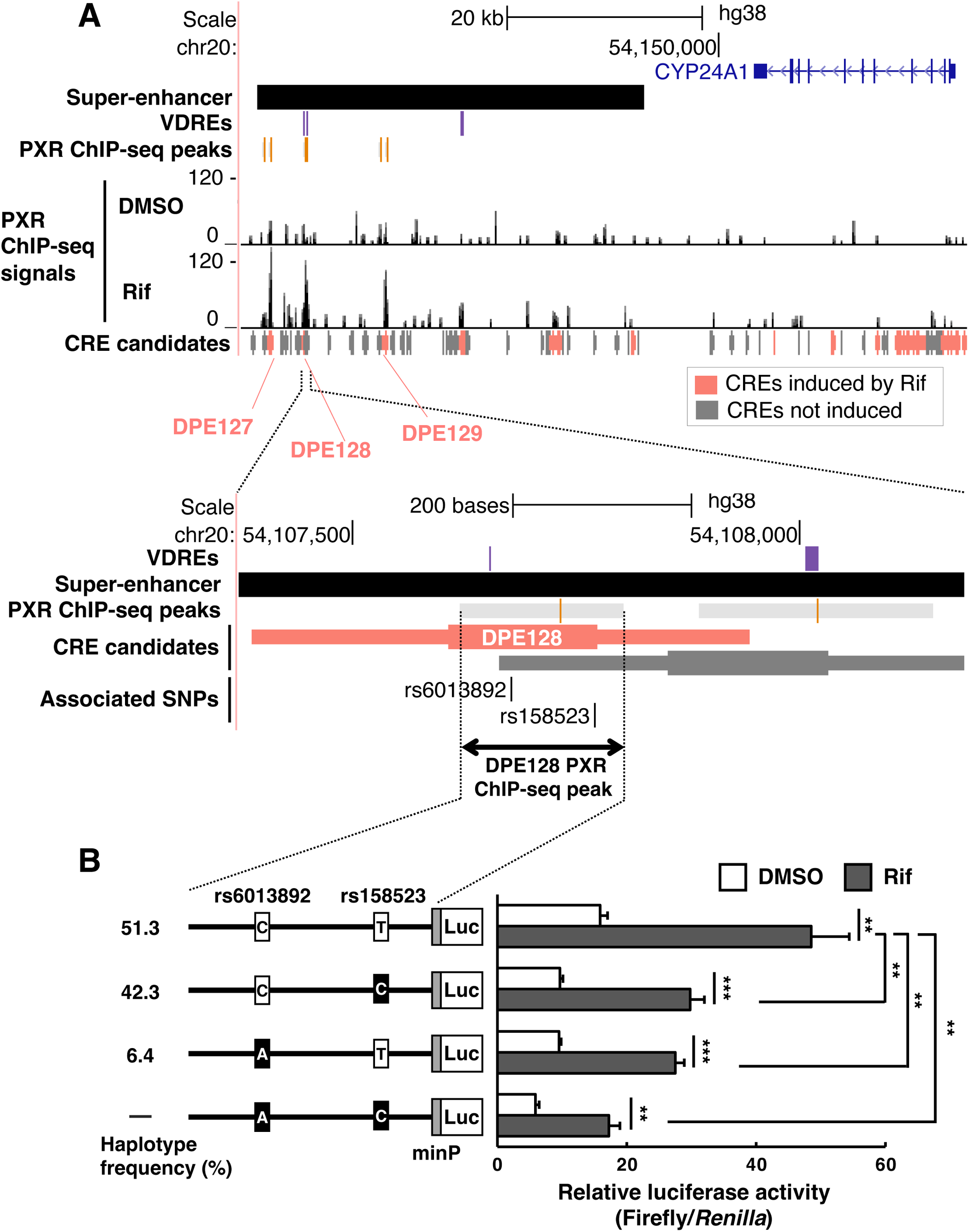
Identification of a drug-inducible downstream super-enhancer of *CYP24A1*. **A** A view of the *CYP24A1* locus and its downstream region in UCSC Genome Browser. Pregnane X receptor (PXR) ChIP-seq peaks in rifampicin (Rif)-treated primary hepatocytes and the *cis*-regulatory element (CRE) candidates are shown aligned with a super-enhancer and previously reported VDREs [53]. ChIP-seq signals indicate normalized read coverage per 50 bp bin in reads per kilobase per million mapped reads. **B** Luciferase activity of DPE128 PXR ChIP-seq peak (+66,190/+66,358 from *CYP24A1* transcription start site) with the minimal promoter (minP) in rifampicin (Rif)-treated ShP51 cells. Three major haplotypes were assessed, and firefly luciferase activity was normalized to *Renilla* luciferase activity. Results are expressed as fold change compared with an empty vector control. The experiment was performed in quadruplicate for each condition and repeated four times with similar results. Representative result was shown. The error bars indicate standard deviations, and unpaired Welch’s *t*-test was used to calculate the *P* values. \*\**P* < 0.01, and \*\*\**P* < 0.001.

**Fig. 4.**
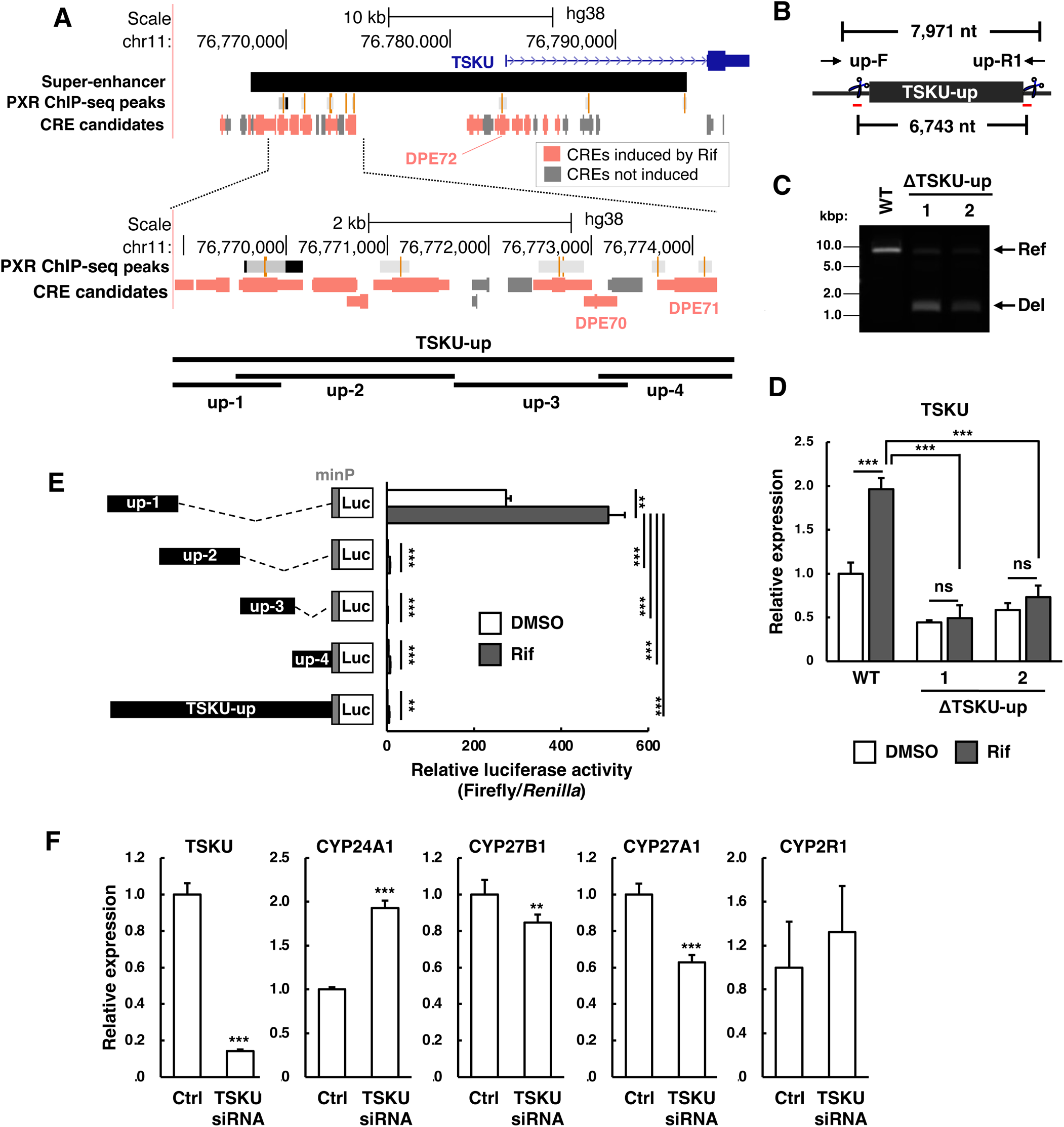
Identification of a drug-inducible super-enhancer of *TSKU*. **A** A view of the *TSKU* locus and its upstream region in UCSC Genome Browser. A zoom-in view indicates the region from 9 kb to 15 kb upstream of *TSKU* (TSKU-up: chr11:76,767,944–76,774,353). TSKU-up was divided into four regions (up-1, −2, −3, and −4) to analyze their transcriptional activity in a luciferase reporter assay. **B** Schematic view of CRISPR/Cas9-based deletion of TSKU-up in ShP51 cells with PCR primers used for amplification of genomic DNA. nt, nucleotides. **C** Gel images show the PCR products from genomic DNA extracted from the deletion mutants (ΔTSKU-up 1, 2). **D** qRT-PCR analysis of *TSKU* mRNA in rifampicin (Rif)-treated ΔTSKU-up 1, 2. *TSKU* mRNA levels were normalized to *GAPDH* mRNA levels. **E** Luciferase activity of each of up-1, −2, −3, −4 and TSKU-up with the minimal promoter (minP) in Rif-treated ShP51 cells. Firefly luciferase activity was normalized to *Renilla* luciferase activity. Results are expressed as fold change compared with an empty vector control. **F** qRT-PCR analysis of *TSKU*, *CYP24A1*, *CYP27B1*, *CYP27A1*, and *CYP2R1* mRNA in *TSKU* siRNA-transfected ShP51 cells. mRNA levels were normalized to *GAPDH* mRNA levels. All experiments were performed in triplicate or quadruplicate for each condition and repeated at least three times with similar results. Representative data were shown. The error bars indicate standard deviations, and unpaired Welch’s *t*-test was used to calculate the *P* values. \*\**P* < 0.01, and \*\*\**P* < 0.001; ns, not significant.

### Functional copy of the well-known enhancer of *UGT1A1*

DPE15 and DPE17 are novel PXR-bound enhancer candidates in the *UGT1A1* locus (Fig. 2A), and we unexpectedly found that the genome sequence of DPE15 is 92% identical to that of PBREM (Fig. 2B). DPE15 (referred to hereafter as PBREM-like) is located at 14.9 kb upstream of the *UGT1A1* promoter, whereas PBREM is located at 3 kb upstream (Fig. 2A). PXR response elements within PBREM (direct repeat (DR)-4, PXRE, gtNR-1, and DR-3 [20]) were mostly preserved in PBREM-like, except that DR-3 had a single nucleotide mismatch to that in PBREM (Fig. 2B). Notably, a SNP (rs4124874, UGT1A1*60 T>G) has been reported in the position of the mismatch, and the minor allele has been shown to lead to a decrease in regulatory activity [21].

To investigate the regulatory role of PBREM-like, we generated homozygous deletion mutants of ShP51 cells using the CRISPR/Cas9 system. We confirmed that homozygous deletion of PBREM (ΔPBREM 1 and 2; Fig. 2C, E) significantly decreased both basal and drug-induced expression of *UGT1A1* (Fig. 2G), consistent with a previous study [20]. Homozygous deletion of PBREM-like (ΔPBREM-like 1 and 2; Fig. 2D, F) showed a significant reduction of *UGT1A1* expression as well (Fig. 2G).

Our luciferase assays found comparable regulatory activities of PBREM-like with the minor alleles of PBREM (Fig. S2A). Our results demonstrate that PBREM-like play a role of enhancer of *UGT1A1* as a functional copy of PBREM, beyond its sequence similarity to the well-known enhancer.

### Intronic enhancer of *UGT1A1*

We subsequently investigated the functional role of DPE17, an enhancer candidate located in the first intron of *UGT1A1*. We found that homozygous deletion (ΔIntronic enh. 1–3; Fig. 2H, I) substantially decreased both basal and drug-induced expression of *UGT1A1* (Fig. 2J). We further confirmed that its silencing by CRISPR interference (CRISPRi) similarly decreased the expression of *UGT1A1* (Fig. S2B).

We focused on two SNPs associated with bilirubin levels (rs3771341 [37, 38] and rs4148325 [38–42]) overlapping with the intronic enhancer (Table 1), and additional two SNPs in the proximal region (rs6742078 [37, 38, 43–45] and rs4148324 [38]). They comprise three haplotypes (Fig. 2K) in the whole population [46, 47] and all four SNPs were significantly associated with the expression level of *UGT1A1* in liver according to the database of GTEx eQTL [48] with a *P* value < 1e−5 and effect size −0.3 (Table S3). We constructed reporter vectors containing a minimal promoter and the +3,657/+4,741 region from *UGT1A1* transcription start site, corresponding to the three haplotypes (Fig. 2K), and found a significant decrease of luciferase activity in both of the two minor alleles. The results identified a novel enhancer of *UGT1A1* with revealing its haplotypes causes alteration of the expression level.

### Impact of *UGT1A1* enhancers on its alternative promoters

The *UGT1A* locus contain nine alternative promoters producing distinct first exons followed by four shared exons [49]. Among these, five isoforms other than *UGT1A1*, namely *UGT1A3*, *UGT1A4*, *UGT1A6*, *UGT1A7*, and *UGT1A9,* have been shown to be expressed in human liver, human primary hepatocytes, and HepG2 cells [50, 51], which is also found in our CAGE data (Fig. S3A). We asked whether the known and the newly identified enhancers function equivalently as enhancers for these promoters producing distinct isoforms.

We examined the expression levels of the *UGT1A1* isoforms in the deletion mutants of PBREM, PBREM-like, and the intronic enhancer (Fig. 2L; Fig. S3B, C). Deletion of the intronic enhancer reduced the expression of all five *UGT1A* isoforms, while deletion of PBREM increased expression of the *UGT1A* isoforms, except for *UGT1A1* and *UGT1A3* (Fig. S3B, C). The deletion effects of PBREM-like were inconclusive due to inconsistencies across the replicates (Fig. S3B). These results indicated that (i) the intronic enhancer act as an enhancer for all *UGT1A* isoforms, (ii) PBREM contributes as enhancer to the two most downstream promoters producing *UGT1A1* and *UGT1A3*, while inhibiting the other *UGT1A* isoforms, and (iii) PBREM-like specifically targets the *UGT1A1* promoter. The effects of the three enhancers in the *UGT1A* locus vary depending on the target promoters (Fig. 2L).

### Variants in a drug-induced super-enhancer alter *CYP24A1* expression

Vitamin D deficiency by PXR activators is likely explained by upregulation of *CYP24A1*, which encodes an enzyme facilitating excretion of vitamin D [52]. Although PXR binds the *CYP24A1* promoter in the human hepatocellular carcinoma Huh7 cell line that overexpress human PXR [52], we found no peaks of PXR ChIP-seq within the *CYP24A1* promoter in human primary hepatocytes (Fig. 3A). This indicates that PXR involvement in *CYP24A1* induction is mediated primarily by distal elements rather than the promoter *in vivo*.

Of the drug-inducible and PXR-bound enhancer candidates, three (DPE127–129) are located downstream of *CYP24A1* (Fig. 3A). They comprise a part of *CYP24A1* enhancer clusters activated by another nuclear receptor, vitamin D receptor (VDR, or NR1I1), through vitamin D response elements (VDREs) [53]. The enhancer cluster overlaps a super-enhancer defined on the basis of H3K27ac ChIP-seq according to the Super-Enhancer database [54]. Binding of PXR to VDRE is possible as PXR recognize similar DNA motifs to VDR [55]. Notably, the region contains rs6013892, a GWAS SNP associated with vitamin D levels [56], and rs158523, genetically correlated (*R*^2^ > 0.8) with another GWAS SNP associated with vitamin D levels, rs158528 [57]. We asked whether the two variants alter the PXR-mediated regulatory activities.

To address the hypothesis, we performed a luciferase reporter assay in ShP51 cells. The major haplotype (rs6013892-C and rs158523-T) had the strongest transcriptional activity at both basal and drug-induced levels (Fig. 3B). The activity was significantly reduced when either of the SNPs was altered and was lowest when both SNPs were altered (Fig. 3B). The results identified that the two SNPs have causal effect to alter the regulatory activity.

### Drug-inducible super-enhancer of *TSKU* alters the expression of vitamin D–metabolizing enzymes

Of the enhancer candidates, three (DPE70–72) were identified at 10.2 kb, 9.4 kb, and 0.5 kb upstream of *TSKU*. PXR was shown to contribute to an increase of *TSKU* expression in phenobarbital-treated human primary hepatocytes [58], which is also found in our CAGE data (Fig. 4A). A series of drug-inducible CREs were found from 8 kb to 13 kb upstream of *TSKU* (Fig. 4A), which is included in a super-enhancer according to the Super-Enhancer database [54]. We generated two clones of ShP51 cells with homozygous deletion of the 6.7 kbp region (ΔTSKU-up 1 and 2; Fig. 4A, B, C; Fig. S4A). We found a significant decrease of *TSKU* expression in the deletion mutants (Fig. 4D), which indicated that a portion of the large region play a role to increases *TSKU* expression. We selected four segments of the region (up-1, −2, −3, and −4) (Fig. 4A) and assessed their regulatory activity with a luciferase reporter assay (Fig. 4E). Although rifampicin increased the activity of all of them, up-1 had the strongest activity in both basal and drug-induced conditions (Fig. 4E; Fig. S4B). These results suggest that up-1 is the main contributor in increasing *TSKU* expression level. Because up-4 contains a SNP rs1149605 that is genetically associated with vitamin D levels [56] [59] (Fig. S4C), we assessed the regulatory activity of the region with a luciferase reporter assay and found no allele-dependency (Fig. S4D). This is consistent with the result that up-1 is the main component within the super-enhancer.

We investigated whether *TSKU* itself affect expression of genes involved in the metabolism of vitamin D. Using siRNA, we reduced the *TSKU* mRNA level to approximately 15% of the original level in ShP51 cells (Fig. 4F). Unexpectedly, this reduction led to an increase of *CYP24A1* expression, which facilitate excretion of vitamin D, and decreases of *CYP27A1* and *CYP27B1* expression, while no effect on *CYP2R1* (Fig. 4F). Given the three latter genes are involved in producing the active form of vitamin D, our results suggested that *TSKU* upregulates the levels of vitamin D by repression of negatively impacting genes (*CYP24A1*) and activation of positively impacting genes (*CYP27A1* and *CYP27B1*).

## Discussion

In this study, we systematically analyzed drug-inducible CREs to elucidate the genomic basis of both anticipated and unanticipated drug responses. We first explored potential CREs through a quantitative and genome-wide assessment of the transcription initiations by using CAGE. The resulting candidates were substantially enriched in loci genetically associated with several biochemical markers measured in serum. Integrative analysis with a reprocessed dataset of PXR ChIP-seq from human primary hepatocytes led us to identify 103 drug-inducible, PXR binding, and GWAS-associated CREs. Targeted experiments, including CRISPR/Cas9 knockout, luciferase reporter assay, and siRNA knockdown provided new insights into PXR downstream regulation. It included as novel enhancers and super-enhancers, allele dependency of the regulatory activities, and an unexpected regulatory cascade leading to ADR. Despite extensive research on drug-responses over several decades, our study identified novel regulatory interactions, made possible by the greater sensitivity of our quantitative transcriptome-based approach (Fig. 5).

**Fig. 5.**
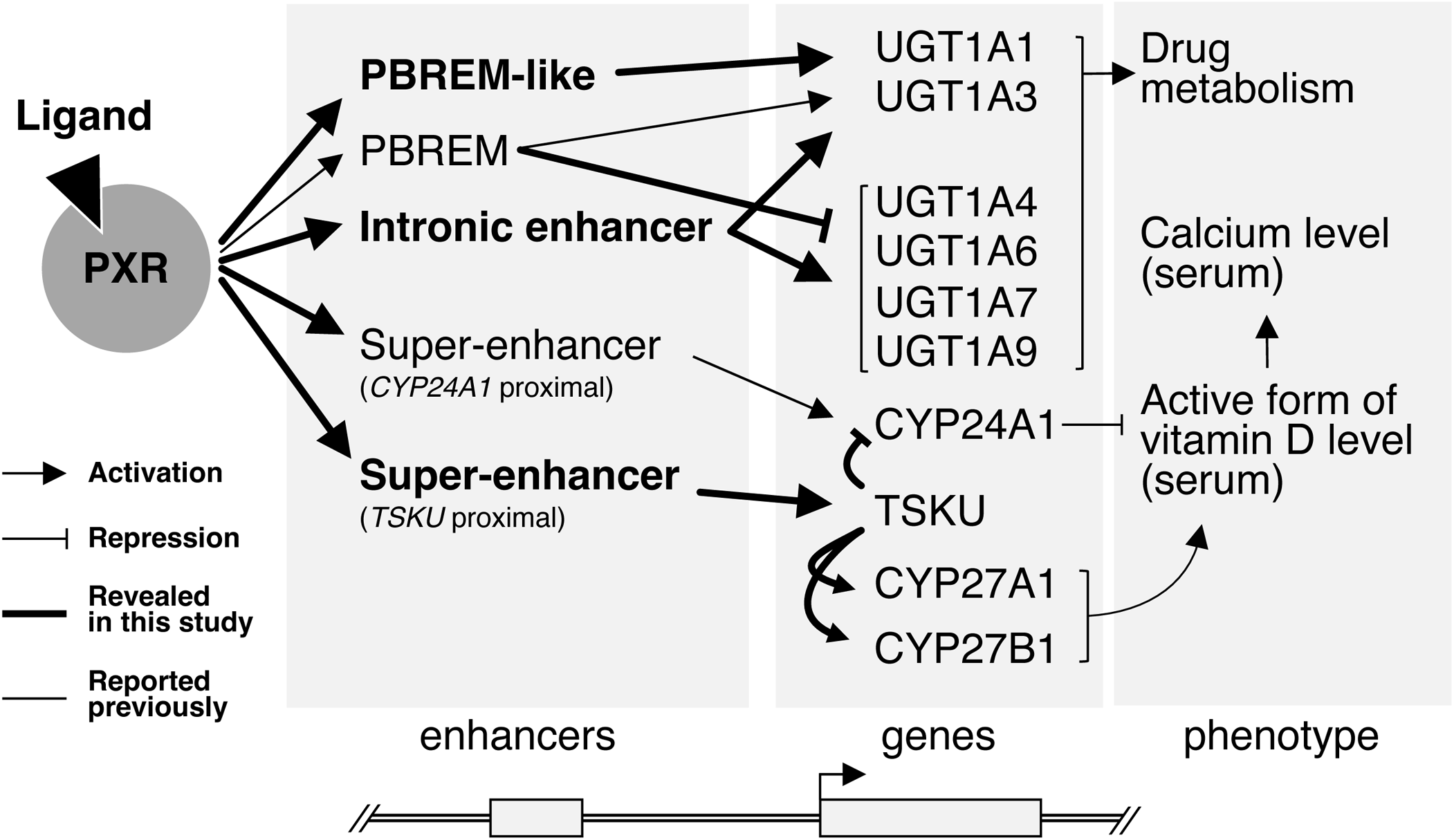
Drug-inducible and pregnane X receptor (PXR)-mediated enhancers and super-enhancers at the *UGT1A1*, *CYP24A1*, and *TSKU* loci. The novel and known enhancers, their regulatory targets, and the resulting phenotypes are illustrated schematically.

### Regulatory elements and alleles associated with drug efficacy

*UGT1A1* plays a critical role in detoxification of a variety of xenobiotics, including therapeutic drugs, and metabolic clearance of the endogenous toxin bilirubin. Whereas PBREM has been the only well-studied enhancer of the gene, we discovered two additional enhancers, PBREM-like and the intronic enhancer, both of which substantially affect *UGT1A1* expression levels.

PBREM-like had 92% sequence identity to PBREM, which is also found in rhesus macaque but not in common marmoset (Fig. S5). This suggests that PBREM-like originated from a segmental duplication of the genome after the divergence of old world monkeys and new world monkeys. Both of the enhancers are crucial for ensuring proper *UGT1A1* expression levels, as demonstrated by the significant decrease of expression observed in the knockout cells (Fig. 2G). The segmental duplication event may have contributed to the very high level of expression of *UGT1A1*. The identification of the intronic enhancer as a novel functional component is both intriguing and important for understanding genotype-dependent responses to drugs. Variants of this enhancer result in substantially lower regulatory activity (Fig. 2K).

The *UGT1A* locus contains nine active CREs in human hepatocytes, including six promoters corresponding to distinct isoforms (UGT1A1, UGT1A3, UGT1A4, UGT1A6, UGT1A7, and UGT1A9), and three enhancers (PBREM, PBREM-like, and the intronic enhancer). Our results indicate that the interactions among these elements can have both positive and negative effects. The impact of allele differences will be influenced by the interplay among the multiple CREs (Fig. 2L). Therefore, a composite model would be required to accurately estimate regulatory outcomes.

### ADR-associated regulatory interactions mediated by super-enhancers

Activation of PXR leads to the upregulation of *CYP24A1*, which encodes an enzyme facilitating excretion of vitamin D [52]. This upregulation may contribute to vitamin D deficiency, an ADR of PXR activators. We found several CREs induced by PXR that included polymorphisms genetically associated with vitamin D levels. The CREs were located within two super-enhancers near *CYP24A1* and *TSKU* loci (Fig. 3A, Fig. 4A).

In contrast to an earlier observation of PXR binding to the *CYP24A1* promoter in a tumor cell line [52], the reprocessed ChIP-seq data obtained from human primary hepatocytes revealed PXR binding within the super-enhancer region rather than the promoter (Fig. 3A). This suggests that PXR-mediated activation of *CYP24A1 in vivo* is primarily mediated by the super-enhancer. We found that two SNPs genetically associated with vitamin D levels actually alter the regulatory activity, supporting the contribution of the super-enhancer to drug responses (Fig. 3B).

The second super-enhancer was located upstream of *TSKU*, which encodes a hepatokine likely involved in energy expenditure [60]. We found that the drug-inducible super-enhancer contributed to the upregulation of *TSKU* (Fig. 4D). Notably, our siRNA experiment revealed that TSKU upregulates the levels of vitamin D by repression of *CYP24A1* gene and activation of *CYP27A1* and *CYP27B1* genes (Fig. 4F). Because *TSKU* is secreted from hepatocytes, this suppression of *CYP24A1* might happen in cells other than hepatocytes and thus may contribute to fine-tuning *CYP24A1* expression by coordinating with other regulatory elements, including the super-enhancer adjacent to *CYP24A1*.

## Conclusions

Our genome-wide quantitative assessment revealed a number of drug-inducible, PXR-mediated, and trait-associated enhancers in human hepatocytes. The results not only aligned closely with the anticipated outcomes to drug exposure but also provided insights into the complex regulatory mechanisms underlying drug responses. The novel enhancers, regulatory alleles, and a molecular cascade leading to ADR contribute a precise understanding of the noncoding elements of the human genome underlying drug responses.

## Methods

### Chemicals and reagents

Restriction enzyme *Sac*I-HF was purchased from New England Biolabs, NEB (Ipswich, MA, USA) and luciferase reporter vectors, pGL4.26 and pGL4.74, from Promega (Madison, WI, USA). The oligonucleotide primers were custom-synthesized by FASMAC Co., Ltd. (Kanagawa, Japan).

### Cell culture

The human hepatocellular carcinoma–derived HepG2 cells that stably express human PXR (ShP51 cells) were kindly provided by Dr. Masahiko Negishi [31]. ShP51 cells and CRISPR/Cas9-mediated enhancer-deleted ShP51 cell lines were cultured in Dulbecco’s modified Eagle’s medium (high glucose) supplemented with 10% fetal bovine serum (Thermo Fisher Scientific, Waltham, MA, USA), 1× MEM Non-Essential Amino Acids Solution (Thermo Fisher Scientific), 100 U/mL penicillin, and 100 μg/mL streptomycin (Thermo Fisher Scientific). Cells were maintained in a humidified incubator, which provided an atmosphere of 5% CO_2_ and 95% air at a constant temperature of 37°C.

### Cell treatment

Cells were treated with dimethyl sulfoxide (DMSO), or with 10 µM or 100 µM rifampicin dissolved in DMSO (Sigma-Aldrich, Burlington, MA, USA) for 24 h.

### Plasmid construction

To construct pGL4.26-1A1-Intronic enh. (Ref, Alt1, or Alt2), the *UGT1A1* intronic region (+3,658/+4,757 from the *UGT1A1* TSS) was amplified by nested polymerase chain reaction (PCR). The amplified DNA fragments were purified and cloned into the *Sac*I-digested pGL4.26 vector. To construct pGL4.26-1A1-Intronic enh. with deleted PXR ChIP-seq peak(Δpeak), the *UGT1A1* intronic regions (fragment 1: +3,658/+4,003 and fragment 2: +4,307/+4,757, both the from the *UGT1A1* TSS) were amplified using pGL4.26-1A1 Intronic enh. (Ref) as a template. The two amplified DNA fragments were cloned into the *Sac*I-digested pGL4.26. To construct pGL4.26-TSKU upstream, 5 upstream regions of *TSKU* (up-1: −15,412/−13,367; up-2: −13,970/−11,658; up-3: −11,677/−10,067; up-4: −10,236/−9,003; and TSKU-up: −15,412/−9,003 from *TSKU* TSS) were amplified by nested PCR. Each of the amplified DNA fragments was cloned into the *Sac*I-digested pGL4.26. Mixed human genomic DNA of 10 people (Promega) was used as a template for 1^st^ round PCR amplification with PrimeSTAR MAX DNA Polymerase (Takara Bio, Shiga, Japan) in a ProFlex PCR System (Thermo Fisher Scientific). The primer sets designed and used in this study are listed in Table S4. To construct pGL4.26-DPE128-PXR ChIP-seq peak (+66,190/+66,358 from *CYP24A1* TSS), the double-stranded DNA fragments containing rs6013892 and rs158523 with different alleles were synthesized (gBlocks; IDT, San Jose, CA, USA). To construct pGL4.26-DPE15 (−14,848/−14,415 from *UGT1A1* TSS) and pGL4.26-DPE16(−3,490/−3,060 from *UGT1A1* TSS), the double-stranded DNA fragments containing rs28900379 and rs11695484 or rs4124874 and rs10929302 with different alleles were synthesized. Each of the gBlocks was cloned into the *Sac*I-digested pGL4.26 vector. The sequences of gBlocks (reference sequences) are listed in Table S5.

All cloning reactions were performed with NEBuilder HiFi DNA Assembly Master Mix (NEB). The constructs were verified by DNA sequencing (Eurofins Scientific, Luxembourg City, Luxembourg).

### Luciferase reporter assay

Cells were transiently reverse-transfected with the constructed pGL4.26 luciferase reporter vector and pGL4.74 vector using Lipofectamine 3000 reagent (Thermo Fisher Scientific). Twenty-four hours after transfection, cells were treated with 10 μM rifampicin for 24 h. Subsequently, cells were resuspended in a passive lysis buffer, and luciferase activity was measured with an ARVOX3 Perkin Elmer 2030 multiple reader (Perkin Elmer, Waltham, MA, USA) using the Dual-Luciferase reporter assay system (Promega). Data were normalized against the *Renilla reniformis* luciferase activity, which served as an internal control.

### Design of single-guide RNAs (sgRNAs)

CHOPCHOP (https://chopchop.cbu.uib.no/) [61], Benchling (https://www.benchling.com/), and CRISPRtarget (http://crispr.otago.ac.nz/CRISPRTarget/crispr_analysis.html) [62] were used to design high-performance gRNAs for the deletion of DPE15, DPE16, and DPE17 around *UGT1A1* and the deletion of TSKU-up. For CRISPRi-mediated repression of the *UGT1A1* intronic region, two gRNAs were designed using CHOPCHOP. The cloning of designed gRNAs into Mammalian Dual-gRNA Expression Lentiviral Vector (pLV-1A1intronic gRNAs) was conducted by Vector Builder (Chicago, IL, USA). All sgRNAs (Alt-R CRISPR-Cas9 sgRNA) for CRISPR/Cas9-mediated deletion were obtained from IDT and are listed in Table S6.

### Transfection and generation of CRISPR/Cas9-mediated enhancer-deletion mutants

Ribonucleoprotein complexes composed of the designed sgRNAs and Alt-R S.p. Cas9 Nuclease V3 (IDT) were electroporated using the NEON transfection system (Thermo Fisher Scientific) into ShP51 cells. Following electroporation, cells were seeded into 10-cm dishes and cultured for 13 days, single-cell clones were hand-picked, grown, and expanded. Genomic DNAs of single-cell clones were extracted using a Wizard SV Genomic DNA Purification Kit (Promega). Deletion was confirmed by genotyping PCR to amplify the targeted region with Tks Gflex DNA Polymerase (Takara Bio); the primers are listed in Table S7. The PCR products were loaded on 0.8% agarose gel containing ethidium bromide and visualized under ultraviolet light. The truncated genomic fragments (mutant PCR products) were subsequently purified and analyzed by DNA sequencing (Eurofins).

### CRISPRi for *UGT1A1* intronic enhancer (DPE17)

pLV-1A1 intronic gRNAs or pLVLscrambled sgRNAs were co-transfected with the ViraPower Lentiviral Packaging Mix into 293FT packaging cells (Thermo Fisher Scientific) by Lipofectamine 3000 reagent (Thermo Fisher Scientific). Viral supernatant was collected and filtered through a 0.45-μm PVDF syringe filter. Viral titration was conducted using a Lenti-X p24 Rapid Titer Kit (Takara Bio). Lentiviral dCas-KRAB or dCas particles, purchased from Vector Builder, were used to infect the recipient ShP51 cells. Forty-eight hours after lentiviral infection, cells were cultured with puromycin (0.25 μg/mL) and hygromycin (400 μg/mL) for 10 days. Subsequently, the transduced cells were processed for quantitative real-time PCR (qRT-PCR).

### siRNA knockdown of *TSKU*

ShP51 cells were transfected with control siRNA (Silencer Select Negative Control #1) or human TSKU siRNA (Silencer Select standard siRNA s24880) from Thermo Fisher Scientific using Lipofectamine RNAiMAX (Thermo Fisher Scientific); 72 h after transfection, cells were processed for qRT-PCR.

### qRT-PCR

Total RNA was isolated using TRIzol reagent (Thermo Fisher Scientific), and quantified by using a Denovix DS-11 spectrophotometer (DeNovix Inc., Wilmington, DE, USA). DNA was digested with TURBO DNase (Thermo Fisher Scientific), and total RNA was reverse-transcribed to cDNA using SuperScript III Reverse Transcriptase (Thermo Fisher Scientific). RT-PCR was performed on CFX96 Real-Time PCR Detection Systems (Bio-Rad, Hercules, CA, USA) using KAPA SYBR FAST qPCR Kit (NIPPON Genetics, Tokyo, Japan). The list of specific sense and antisense oligonucleotide primers is shown in Table S8.

### Statistical analysis

The experiments of luciferase assay and qRT-PCR were performed in triplicate or quadruplicate for each condition and repeated at least three times with similar results. Representative data were shown. Statistical significance was determined using unpaired Welch’s *t*-test, and statistical analyses used the GraphPad Prism5 software (GraphPad Software, San Diego, CA). Data are presented as mean ± standard deviation. The statistical parameters (ns, not significant; \**P* < 0.05; \*\**P* < 0.01; and \*\*\**P* < 0.001) are presented in individual figures and figure legends. A value of *P* < 0.05 was considered statistically significant.

### CAGE and data analysis

ShP51 cells were treated with 10 μM or 100 μM rifampicin for 24 h. Total RNA was isolated using a miRNeasy Mini Kit (Qiagen, Venlo, Netherlands). The quality of total RNA was assessed by an Agilent 2100 Bioanalyzer (Agilent Technologies, Santa Clara, CA, USA) to ensure that the RNA integrity number was over 7.0. CAGE libraries were prepared using a CAGE Library preparation kit (DNAform, Kanagawa, Japan). Briefly, cDNA was synthesized from total RNA using random primers. The ribose diols in the 5′ cap structures of RNA were oxidized and then biotinylated. The biotinylated RNA/cDNAs were selected by streptavidin beads (a process known as cap-trapping). RNA was digested with RNaseONE/H, adaptors were ligated to both ends of cDNA, and CAGE libraries of double-stranded cDNA were constructed. The CAGE libraries were validated for quality and size distribution using an Agilent 2100 Bioanalyzer and sequenced to the sequencing depth of approximately 54 million reads on average by NextSeq500 (Illumina, San Diego, CA, USA).

Obtained reads were aligned with the reference human genome GRCh38 by using STAR (v2.7.5a) [63], taking splice junctions of Gencode transcripts (v37) [64] to improve the alignment accuracy. The first position of each uniquely mapped read was counted as the transcription initiation, and the result was used to identify divergently transcribed regions as candidate CREs by using CREate (https://github.com/hkawaji/CREate), bedtools (v2.29.2) [65], and the utility programs of the UCSC Genome Browser Database (v425) [66]. Read counts starting from the identified candidate regions were subjected to quantitative analysis by using edgeR (v3.30.0) [67], consisting of normalization based on the relative log expression method and differential analysis based on the exact test (FDR < 0.1 for statistical significance).

Functional enrichment of the upregulated CRE candidates was assessed by Genomic Regions Enrichment of Annotations Tool (v4.0.4) [68]. Partitioning heritability of S-LDSC was computed by LDSC (v1.0.1) [36] with the genomic coordinates of the CRE candidates converted into GRCh37 by the liftOver tool [66] and 1000 Genomes Project Phase 3 data (https://doi.org/10.5281/zenodo.7768714).

### ChIP-seq data analysis

Data of PXR ChIP-seq performed on human primary hepatocytes [22] (accession numbers: SRR1642056 and SRR1642057 for PXR ChIP-seq and SRR1642055 for the control experiment) were downloaded and aligned with the reference human genome GRCh38 by BWA MEM (v0.7.17) [69]. Alignments with a mapping quality score more than 20 were subjected to peak call by MACS2 (v2.2.7.1) with the default setting [70]. With a threshold of FDR of <0.2 for detecting the binding of PXR to XREM, 27,633 peaks for dimethyl sulfoxide and 11,752 peaks for rifampicin treatment were identified.

### Integration with the GWAS catalog database

Genomic coordinates of index SNPs in the GWAS Catalog [6] and common SNPs in The Single Nucleotide Polymorphism Database [71] were downloaded from the UCSC Genome Browser Database [72]. The *R*^2^ measure of linkage disequilibrium between SNPs within 200 kbp was calculated by using plink2 (v2.00a2.3LM) [73] and was based on European population data in the 1000 Genomes Project database.

### Determination of haplotypes and estimation of haplotype frequency

The frequency of each haplotype in all populations was obtained from LDlink (data from the 1000 Genomes Project) via LDhap (https://ldlink.nih.gov/).

## Availability of data and materials

The data were deposited to the Gene Expression Omnibus (GEO) database under accession number GSE272109.

## Acknowledgements

We greatly thank Prof. Negishi at National Institute of Environmental Health Sciences for kindly providing ShP51 cells. Computations in this study were partially performed on the NIG supercomputer at ROIS National Institute of Genetics.

## Authors’ contributions

Conceptualization: S.S., and H.K. Methodology: S.S., and H.K. Computational analyses: H.K. Experimental analyses: S.S., and R.W. Data curation: H.K. Writing of the draft: S.S., and H.K. Figures: S.S., and H.K. The authors have read and approved the final manuscript.

## Ethics declarations

### Ethics approval and consent to participate

Not applicable.

### Consent for publication

Not applicable.

## References

1. Roden DM, McLeod HL, Relling MV, Williams MS, Mensah GA, Peterson JF, Van Driest SL: Pharmacogenomics. Lancet 2019, 394:521–532.

2. Takahashi H, Echizen H: Pharmacogenetics of warfarin elimination and its clinical implications. Clin Pharmacokinet 2001, 40:587–603.

3. Yoon YR, Shon JH, Kim MK, Lim YC, Lee HR, Park JY, Cha IJ, Shin JG: Frequency of cytochrome P450 2C9 mutant alleles in a Korean population. Br J Clin Pharmacol 2001, 51:277–280.

4. Alessandrini M, Asfaha S, Dodgen TM, Warnich L, Pepper MS: Cytochrome P450 pharmacogenetics in African populations. Drug Metab Rev 2013, 45:253–275.

5. Maurano MT, Humbert R, Rynes E, Thurman RE, Haugen E, Wang H, Reynolds AP, Sandstrom R, Qu H, Brody J, et al: Systematic localization of common disease-associated variation in regulatory DNA. Science 2012, 337:1190–1195.

6. Sollis E, Mosaku A, Abid A, Buniello A, Cerezo M, Gil L, Groza T, Gunes O, Hall P, Hayhurst J, et al: The NHGRI-EBI GWAS Catalog: knowledgebase and deposition resource. Nucleic Acids Res 2023, 51:D977–D985.

7. Nasser J, Bergman DT, Fulco CP, Guckelberger P, Doughty BR, Patwardhan TA, Jones TR, Nguyen TH, Ulirsch JC, Lekschas F, et al: Genome-wide enhancer maps link risk variants to disease genes. Nature 2021, 593:238–243.

8. Andersson R, Gebhard C, Miguel-Escalada I, Hoof I, Bornholdt J, Boyd M, Chen Y, Zhao X, Schmidl C, Suzuki T, et al: An atlas of active enhancers across human cell types and tissues. Nature 2014, 507:455–461.

9. Finucane HK, Bulik-Sullivan B, Gusev A, Trynka G, Reshef Y, Loh PR, Anttila V, Xu H, Zang C, Farh K, et al: Partitioning heritability by functional annotation using genome-wide association summary statistics. Nat Genet 2015, 47:1228–1235.

10. Murakawa Y, Yoshihara M, Kawaji H, Nishikawa M, Zayed H, Suzuki H, Fantom C, Hayashizaki Y: Enhanced Identification of Transcriptional Enhancers Provides Mechanistic Insights into Diseases. Trends Genet 2016, 32:76–88.

11. Iyer L, Das S, Janisch L, Wen M, Ramirez J, Karrison T, Fleming GF, Vokes EE, Schilsky RL, Ratain MJ: UGT1A1*28 polymorphism as a determinant of irinotecan disposition and toxicity. Pharmacogenomics J 2002, 2:43–47.

12. Marcuello E, Altes A, Menoyo A, Del Rio E, Gomez-Pardo M, Baiget M: UGT1A1 gene variations and irinotecan treatment in patients with metastatic colorectal cancer. Br J Cancer 2004, 91:678–682.

13. Karas S, Innocenti F: All You Need to Know About UGT1A1 Genetic Testing for Patients Treated With Irinotecan: A Practitioner-Friendly Guide. JCO Oncol Pract 2022, 18:270–277.

14. Grober U, Kisters K: Influence of drugs on vitamin D and calcium metabolism. Dermatoendocrinol 2012, 4:158–166.

15. Hosseinpour F, Ellfolk M, Norlin M, Wikvall K: Phenobarbital suppresses vitamin D3 25-hydroxylase expression: a potential new mechanism for drug-induced osteomalacia. Biochem Biophys Res Commun 2007, 357:603–607.

16. Bertilsson G, Heidrich J, Svensson K, Asman M, Jendeberg L, Sydow-Backman M, Ohlsson R, Postlind H, Blomquist P, Berkenstam A: Identification of a human nuclear receptor defines a new signaling pathway for CYP3A induction. Proc Natl Acad Sci U S A 1998, 95:12208–12213.

17. Kliewer SA, Moore JT, Wade L, Staudinger JL, Watson MA, Jones SA, McKee DD, Oliver BB, Willson TM, Zetterstrom RH, et al: An orphan nuclear receptor activated by pregnanes defines a novel steroid signaling pathway. Cell 1998, 92:73–82.

18. Lehmann JM, McKee DD, Watson MA, Willson TM, Moore JT, Kliewer SA: The human orphan nuclear receptor PXR is activated by compounds that regulate CYP3A4 gene expression and cause drug interactions. J Clin Invest 1998, 102:1016–1023.

19. Goodwin B, Hodgson E, Liddle C: The orphan human pregnane X receptor mediates the transcriptional activation of CYP3A4 by rifampicin through a distal enhancer module. Mol Pharmacol 1999, 56:1329–1339.

20. Sugatani J, Kojima H, Ueda A, Kakizaki S, Yoshinari K, Gong QH, Owens IS, Negishi M, Sueyoshi T: The phenobarbital response enhancer module in the human bilirubin UDP-glucuronosyltransferase UGT1A1 gene and regulation by the nuclear receptor CAR. Hepatology 2001, 33:1232–1238.

21. Tomerak RH, Helal NF, Shaker OG, Yousef MA: Association between the Specific UGT1A1 Promoter Sequence Variant (c-3279T>G) and Unconjugated Neonatal Hyperbilirubinemia. J Trop Pediatr 2016, 62:457–463.

22. Smith RP, Eckalbar WL, Morrissey KM, Luizon MR, Hoffmann TJ, Sun X, Jones SL, Force Aldred S, Ramamoorthy A, Desta Z, et al: Genome-wide discovery of drug-dependent human liver regulatory elements. PLoS Genet 2014, 10:e1004648.

23. Kim TK, Hemberg M, Gray JM, Costa AM, Bear DM, Wu J, Harmin DA, Laptewicz M, Barbara-Haley K, Kuersten S, et al: Widespread transcription at neuronal activity-regulated enhancers. Nature 2010, 465:182–187.

24. Yao L, Liang J, Ozer A, Leung AK, Lis JT, Yu H: A comparison of experimental assays and analytical methods for genome-wide identification of active enhancers. Nat Biotechnol 2022, 40:1056–1065.

25. Franco HL, Nagari A, Malladi VS, Li W, Xi Y, Richardson D, Allton KL, Tanaka K, Li J, Murakami S, et al: Enhancer transcription reveals subtype-specific gene expression programs controlling breast cancer pathogenesis. Genome Res 2018, 28:159–170.

26. Cheng JH, Pan DZ, Tsai ZT, Tsai HK: Genome-wide analysis of enhancer RNA in gene regulation across 12 mouse tissues. Sci Rep 2015, 5:12648.

27. Arner E, Daub CO, Vitting-Seerup K, Andersson R, Lilje B, Drablos F, Lennartsson A, Ronnerblad M, Hrydziuszko O, Vitezic M, et al: Transcribed enhancers lead waves of coordinated transcription in transitioning mammalian cells. Science 2015, 347:1010–1014.

28. Consortium F, the RP, Clst, Forrest AR, Kawaji H, Rehli M, Baillie JK, de Hoon MJ, Haberle V, Lassmann T, et al: A promoter-level mammalian expression atlas. Nature 2014, 507:462–470.

29. Kanamori-Katayama M, Itoh M, Kawaji H, Lassmann T, Katayama S, Kojima M, Bertin N, Kaiho A, Ninomiya N, Daub CO, et al: Unamplified cap analysis of gene expression on a single-molecule sequencer. Genome Res 2011, 21:1150–1159.

30. Morioka MS, Kawaji H, Nishiyori-Sueki H, Murata M, Kojima-Ishiyama M, Carninci P, Itoh M: Cap Analysis of Gene Expression (CAGE): A Quantitative and Genome-Wide Assay of Transcription Start Sites. Methods Mol Biol 2020, 2120:277–301.

31. Kodama S, Negishi M: Pregnane X receptor PXR activates the GADD45beta gene, eliciting the p38 MAPK signal and cell migration. J Biol Chem 2011, 286:3570–3578.

32. Yamazaki H, Shimada T: Progesterone and testosterone hydroxylation by cytochromes P450 2C19, 2C9, and 3A4 in human liver microsomes. Arch Biochem Biophys 1997, 346:161–169.

33. Zhou C, Tabb MM, Nelson EL, Grun F, Verma S, Sadatrafiei A, Lin M, Mallick S, Forman BM, Thummel KE, Blumberg B: Mutual repression between steroid and xenobiotic receptor and NF-kappaB signaling pathways links xenobiotic metabolism and inflammation. J Clin Invest 2006, 116:2280–2289.

34. Zhou C, Verma S, Blumberg B: The steroid and xenobiotic receptor (SXR), beyond xenobiotic metabolism. Nucl Recept Signal 2009, 7:e001.

35. Sudlow C, Gallacher J, Allen N, Beral V, Burton P, Danesh J, Downey P, Elliott P, Green J, Landray M, et al: UK biobank: an open access resource for identifying the causes of a wide range of complex diseases of middle and old age. PLoS Med 2015, 12:e1001779.

36. Bulik-Sullivan BK, Loh PR, Finucane HK, Ripke S, Yang J, Schizophrenia Working Group of the Psychiatric Genomics C, Patterson N, Daly MJ, Price AL, Neale BM: LD Score regression distinguishes confounding from polygenicity in genome-wide association studies. Nat Genet 2015, 47:291–295.

37. Chu X, Liu L, Ye J, Wen Y, Li P, Cheng B, Cheng S, Zhang L, Qi X, Ma M, et al: Insomnia affects the levels of plasma bilirubin and protein metabolism: an observational study and GWGEIS in UK Biobank cohort. Sleep Med 2021, 85:184–190.

38. Wanga V, Venuto C, Morse GD, Acosta EP, Daar ES, Haas DW, Li C, Shepherd BE: Genomewide association study of tenofovir pharmacokinetics and creatinine clearance in AIDS Clinical Trials Group protocol A5202. Pharmacogenet Genomics 2015, 25:450–461.

39. Coltell O, Asensio EM, Sorli JV, Barragan R, Fernandez-Carrion R, Portoles O, Ortega-Azorin C, Martinez-LaCruz R, Gonzalez JI, Zanon-Moreno V, et al: Genome-Wide Association Study (GWAS) on Bilirubin Concentrations in Subjects with Metabolic Syndrome: Sex-Specific GWAS Analysis and Gene-Diet Interactions in a Mediterranean Population. Nutrients 2019, 11.

40. Moore CB, Verma A, Pendergrass S, Verma SS, Johnson DH, Daar ES, Gulick RM, Haubrich R, Robbins GK, Ritchie MD, Haas DW: Phenome-wide Association Study Relating Pretreatment Laboratory Parameters With Human Genetic Variants in AIDS Clinical Trials Group Protocols. Open Forum Infect Dis 2015, 2:ofu113.

41. DiStefano JK, Kingsley C, Craig Wood G, Chu X, Argyropoulos G, Still CD, Done SC, Legendre C, Tembe W, Gerhard GS: Genome-wide analysis of hepatic lipid content in extreme obesity. Acta Diabetol 2015, 52:373–382.

42. Bielinski SJ, Chai HS, Pathak J, Talwalkar JA, Limburg PJ, Gullerud RE, Sicotte H, Klee EW, Ross JL, Kocher JP, et al: Mayo Genome Consortia: a genotype-phenotype resource for genome-wide association studies with an application to the analysis of circulating bilirubin levels. Mayo Clin Proc 2011, 86:606–614.

43. Dai X, Wu C, He Y, Gui L, Zhou L, Guo H, Yuan J, Yang B, Li J, Deng Q, et al: A genome-wide association study for serum bilirubin levels and gene-environment interaction in a Chinese population. Genet Epidemiol 2013, 37:293–300.

44. Kang TW, Kim HJ, Ju H, Kim JH, Jeon YJ, Lee HC, Kim KK, Kim JW, Lee S, Kim JY, et al: Genome-wide association of serum bilirubin levels in Korean population. Hum Mol Genet 2010, 19:3672–3678.

45. Johnson AD, Kavousi M, Smith AV, Chen MH, Dehghan A, Aspelund T, Lin JP, van Duijn CM, Harris TB, Cupples LA, et al: Genome-wide association meta-analysis for total serum bilirubin levels. Hum Mol Genet 2009, 18:2700–2710.

46. Genomes Project C, Abecasis GR, Auton A, Brooks LD, DePristo MA, Durbin RM, Handsaker RE, Kang HM, Marth GT, McVean GA: An integrated map of genetic variation from 1,092 human genomes. Nature 2012, 491:56–65.

47. Machiela MJ, Chanock SJ: LDlink: a web-based application for exploring population-specific haplotype structure and linking correlated alleles of possible functional variants. Bioinformatics 2015, 31:3555–3557.

48. Consortium GT: Human genomics. The Genotype-Tissue Expression (GTEx) pilot analysis: multitissue gene regulation in humans. Science 2015, 348:648–660.

49. Ritter JK, Chen F, Sheen YY, Tran HM, Kimura S, Yeatman MT, Owens IS: A novel complex locus UGT1 encodes human bilirubin, phenol, and other UDP-glucuronosyltransferase isozymes with identical carboxyl termini. J Biol Chem 1992, 267:3257–3261.

50. Nakamura A, Nakajima M, Yamanaka H, Fujiwara R, Yokoi T: Expression of UGT1A and UGT2B mRNA in human normal tissues and various cell lines. Drug Metab Dispos 2008, 36:1461–1464.

51. Senekeo-Effenberger K, Chen S, Brace-Sinnokrak E, Bonzo JA, Yueh MF, Argikar U, Kaeding J, Trottier J, Remmel RP, Ritter JK, et al: Expression of the human UGT1 locus in transgenic mice by 4-chloro-6-(2,3-xylidino)-2-pyrimidinylthioacetic acid (WY-14643) and implications on drug metabolism through peroxisome proliferator-activated receptor alpha activation. Drug Metab Dispos 2007, 35:419–427.

52. Pascussi JM, Robert A, Nguyen M, Walrant-Debray O, Garabedian M, Martin P, Pineau T, Saric J, Navarro F, Maurel P, Vilarem MJ: Possible involvement of pregnane X receptor-enhanced CYP24 expression in drug-induced osteomalacia. J Clin Invest 2005, 115:177–186.

53. Meyer MB, Goetsch PD, Pike JW: A downstream intergenic cluster of regulatory enhancers contributes to the induction of CYP24A1 expression by 1alpha,25-dihydroxyvitamin D3. J Biol Chem 2010, 285:15599–15610.

54. Wang Y, Song C, Zhao J, Zhang Y, Zhao X, Feng C, Zhang G, Zhu J, Wang F, Qian F, et al: SEdb 2.0: a comprehensive super-enhancer database of human and mouse. Nucleic Acids Res 2023, 51:D280–D290.

55. Penvose A, Keenan JL, Bray D, Ramlall V, Siggers T: Comprehensive study of nuclear receptor DNA binding provides a revised framework for understanding receptor specificity. Nat Commun 2019, 10:2514.

56. Revez JA, Lin T, Qiao Z, Xue A, Holtz Y, Zhu Z, Zeng J, Wang H, Sidorenko J, Kemper KE, et al: Genome-wide association study identifies 143 loci associated with 25 hydroxyvitamin D concentration. Nat Commun 2020, 11:1647.

57. Sinnott-Armstrong N, Tanigawa Y, Amar D, Mars N, Benner C, Aguirre M, Venkataraman GR, Wainberg M, Ollila HM, Kiiskinen T, et al: Genetics of 35 blood and urine biomarkers in the UK Biobank. Nat Genet 2021, 53:185–194.

58. de Boussac H, Gondeau C, Briolotti P, Duret C, Treindl F, Romer M, Fabre JM, Herrero A, Ramos J, Maurel P, et al: Epidermal Growth Factor Represses Constitutive Androstane Receptor Expression in Primary Human Hepatocytes and Favors Regulation by Pregnane X Receptor. Drug Metab Dispos 2018, 46:223–236.

59. Manousaki D, Mitchell R, Dudding T, Haworth S, Harroud A, Forgetta V, Shah RL, Luan J, Langenberg C, Timpson NJ, Richards JB: Genome-wide Association Study for Vitamin D Levels Reveals 69 Independent Loci. Am J Hum Genet 2020, 106:327–337.

60. Wang Q, Sharma VP, Shen H, Xiao Y, Zhu Q, Xiong X, Guo L, Jiang L, Ohta K, Li S, et al: The hepatokine Tsukushi gates energy expenditure via brown fat sympathetic innervation. Nat Metab 2019, 1:251–260.

61. Labun K, Montague TG, Krause M, Torres Cleuren YN, Tjeldnes H, Valen E: CHOPCHOP v3: expanding the CRISPR web toolbox beyond genome editing. Nucleic Acids Res 2019, 47:W171–W174.

62. Biswas A, Gagnon JN, Brouns SJ, Fineran PC, Brown CM: CRISPRTarget: bioinformatic prediction and analysis of crRNA targets. RNA Biol 2013, 10:817–827.

63. Dobin A, Davis CA, Schlesinger F, Drenkow J, Zaleski C, Jha S, Batut P, Chaisson M, Gingeras TR: STAR: ultrafast universal RNA-seq aligner. Bioinformatics 2013, 29:15–21.

64. Frankish A, Diekhans M, Jungreis I, Lagarde J, Loveland JE, Mudge JM, Sisu C, Wright JC, Armstrong J, Barnes I, et al: Gencode 2021. Nucleic Acids Res 2021, 49:D916–D923.

65. Quinlan AR, Hall IM: BEDTools: a flexible suite of utilities for comparing genomic features. Bioinformatics 2010, 26:841–842.

66. Kent WJ, Zweig AS, Barber G, Hinrichs AS, Karolchik D: BigWig and BigBed: enabling browsing of large distributed datasets. Bioinformatics 2010, 26:2204–2207.

67. Robinson MD, McCarthy DJ, Smyth GK: edgeR: a Bioconductor package for differential expression analysis of digital gene expression data. Bioinformatics 2010, 26:139–140.

68. McLean CY, Bristor D, Hiller M, Clarke SL, Schaar BT, Lowe CB, Wenger AM, Bejerano G: GREAT improves functional interpretation of cis-regulatory regions. Nat Biotechnol 2010, 28:495–501.

69. Li H: Aligning sequence reads, clone sequences and assembly contigs with BWA-MEM. arXiv: Genomics 2013, arXiv:1303.3997.

70. Zhang Y, Liu T, Meyer CA, Eeckhoute J, Johnson DS, Bernstein BE, Nusbaum C, Myers RM, Brown M, Li W, Liu XS: Model-based analysis of ChIP-Seq (MACS). Genome Biol 2008, 9:R137.

71. Sherry ST, Ward MH, Kholodov M, Baker J, Phan L, Smigielski EM, Sirotkin K: dbSNP: the NCBI database of genetic variation. Nucleic Acids Res 2001, 29:308–311.

72. Nassar LR, Barber GP, Benet-Pages A, Casper J, Clawson H, Diekhans M, Fischer C, Gonzalez JN, Hinrichs AS, Lee BT, et al: The UCSC Genome Browser database: 2023 update. Nucleic Acids Res 2023, 51:D1188–D1195.

73. Chang CC, Chow CC, Tellier LC, Vattikuti S, Purcell SM, Lee JJ: Second-generation PLINK: rising to the challenge of larger and richer datasets. Gigascience 2015, 4:7.

74. Sugatani J, Nishitani S, Yamakawa K, Yoshinari K, Sueyoshi T, Negishi M, Miwa M: Transcriptional regulation of human UGT1A1 gene expression: activated glucocorticoid receptor enhances constitutive androstane receptor/pregnane X receptor-mediated UDP-glucuronosyltransferase 1A1 regulation with glucocorticoid receptor-interacting protein 1. Mol Pharmacol 2005, 67:845–855.

