## Supplemental Figures for "Drug-induced cis-regulatory elements in human hepatocytes affect molecular phenotypes associated with adverse reactions"

Fig. S1

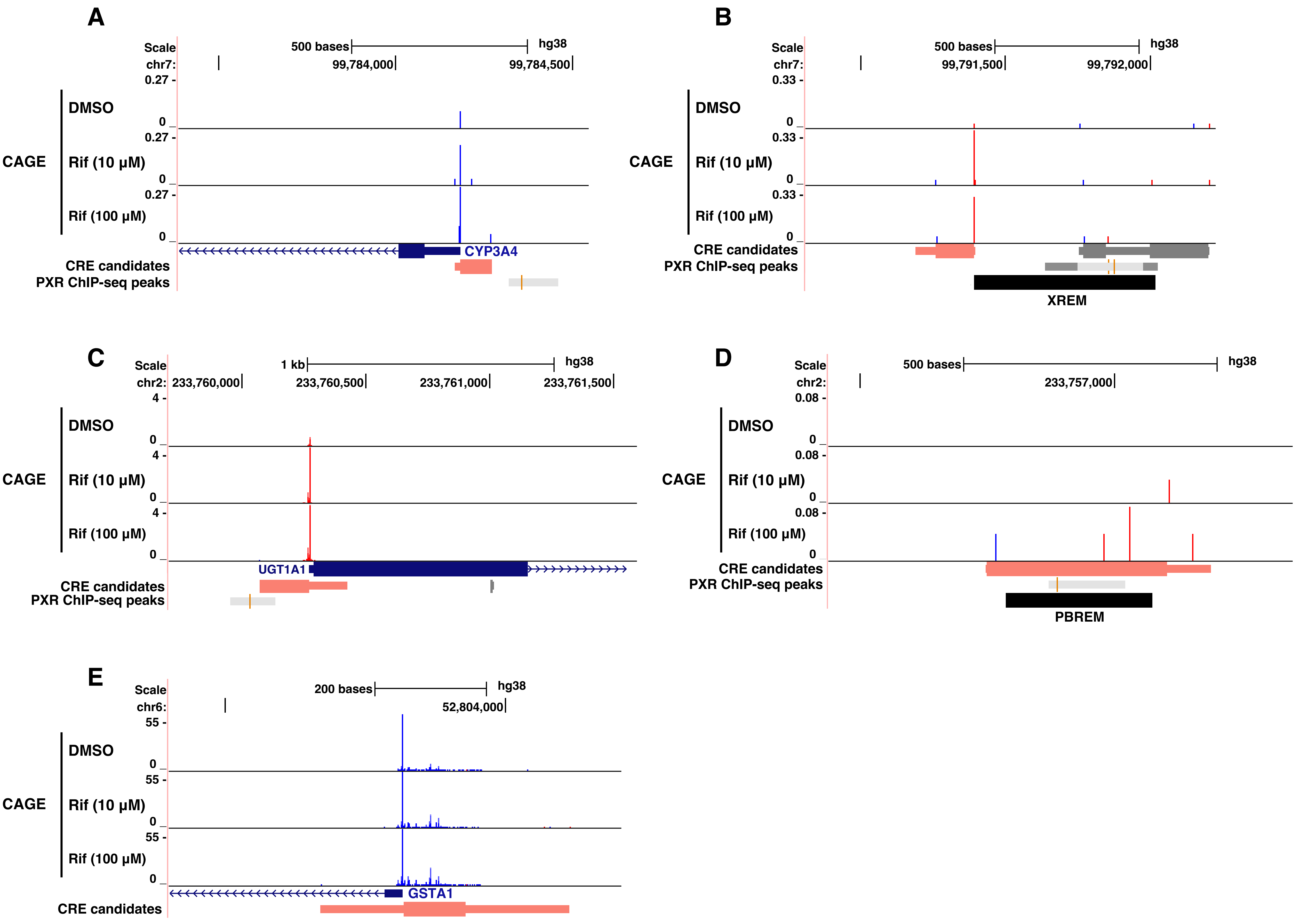


**Fig. S1**
A view of the promoters, *CYP3A4* (A), *UGT1A1* (C), and *GSTA1* (E) and enhancers, XREM (B) and PBREM (D) in UCSC Genome Browser. The tracks for cap analysis of gene expression (CAGE) indicate the frequencies of monitored transcription start sites at single base-pair (bp) resolution in transcripts per million, where red and blue indicate the orientation of transcription (red: forward, blue: reverse).

**Fig. S2**

**A**

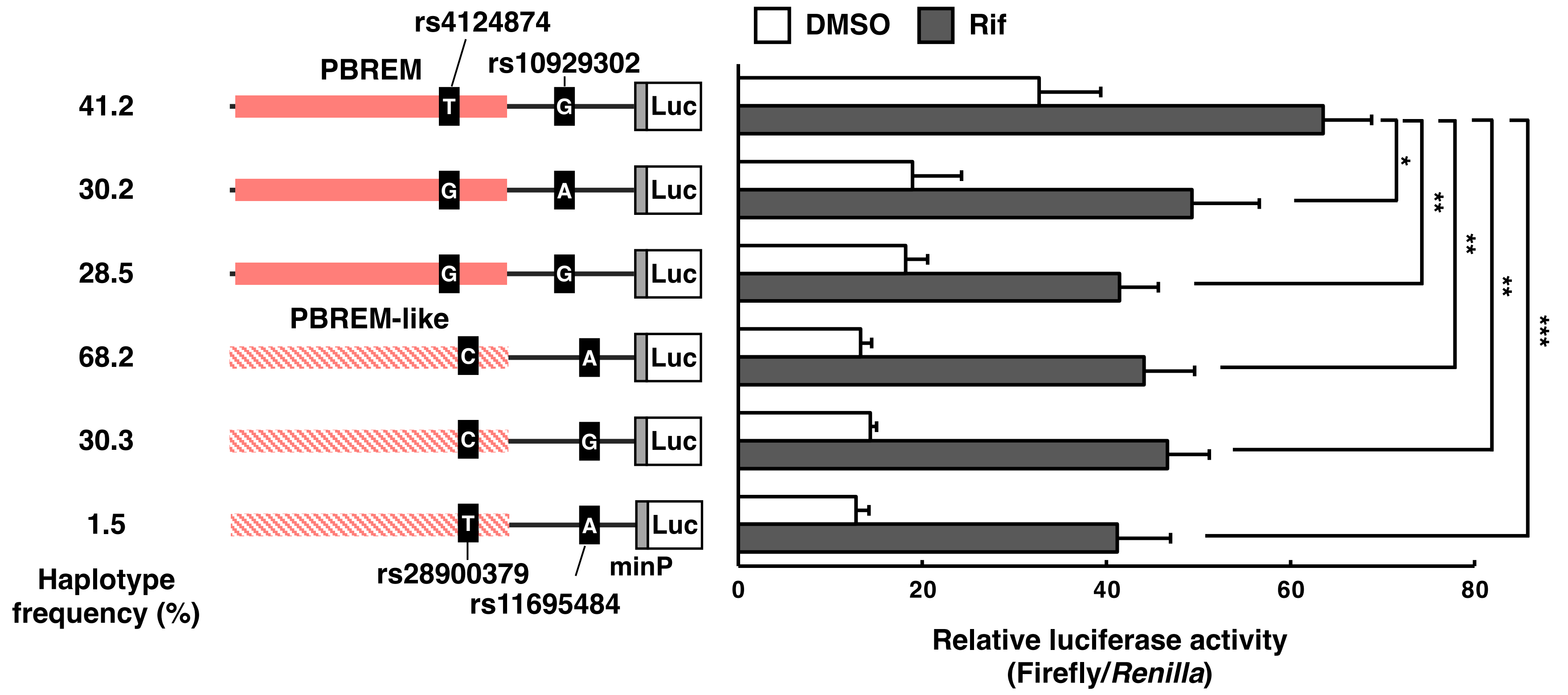


**B**

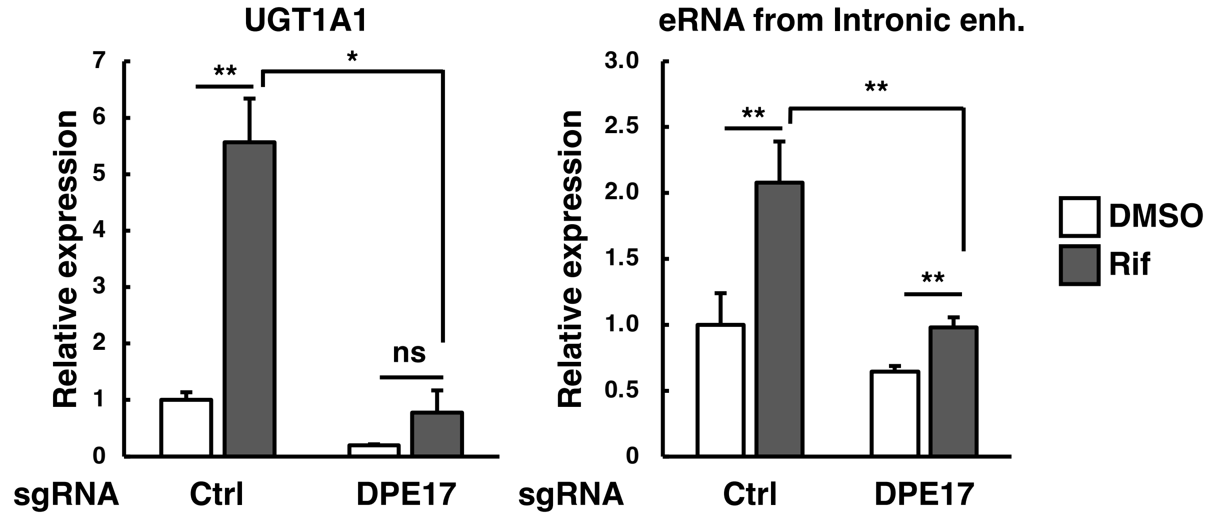


**Fig. S2**

**A** Luciferase activity of -3,490/-3,060 (PBREM, DPE16) region or -14,848/-14,415 (PBREM-like, DPE15) region from *UGT1A1* transcription start site with minimal promoter (minP) in rifampicin (Rif)-treated ShP51 cells. There are three major haplotypes with the frequencies in the population respectively. Firefly luciferase activity was normalized to *Renilla* luciferase activity. Results are shown as fold changes compared with empty vector control. **B** qRT-PCR analysis of *UGT1A1* mRNA and eRNA transcribed from *UGT1A1* Intronic enh. (DPE17) in Rif-treated ShP51 cells which expresses dCas9-KRAB and sgRNAs targeting the DPE17. Both mRNA and eRNA levels are normalized to *GAPDH* mRNA levels. All experiments were performed in triplicate or quadruplicate for each condition and repeated at least three times with similar results. Representative data were shown. The error bars indicate standard deviation, and unpaired Welch's *t*-test was used to calculate the *P* value. *$P < 0.05$, **$P < 0.01$, and ***$P < 0.001$.

Fig. S3

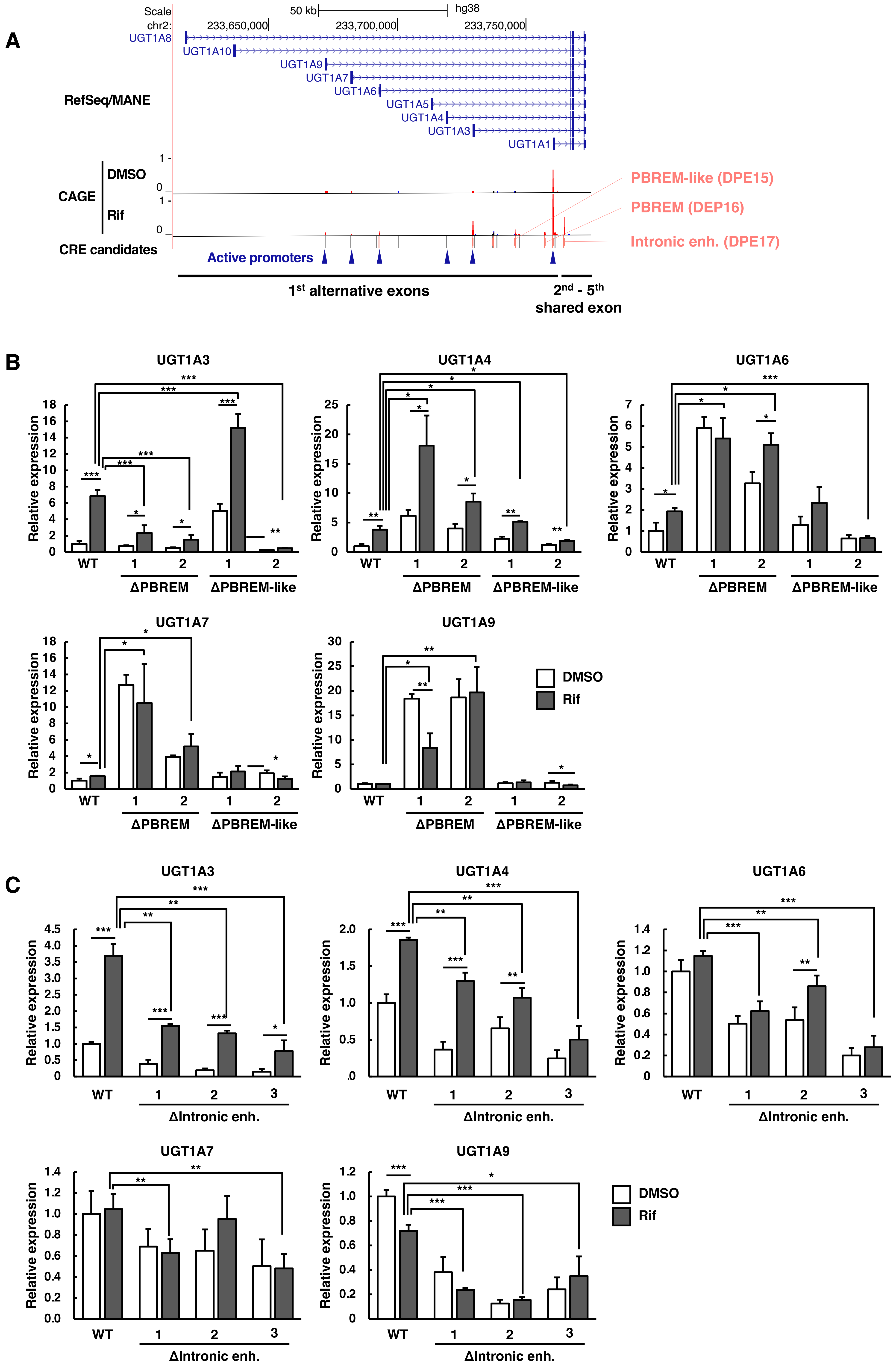


**Fig. S3**
**A** A view of *UGT1A* locus in UCSC Genome Browser. The locus includes unique alternate first exons followed by four common exons. DPE15 (PBREM-like), DPE16 (PBREM ), and DPE17 (Intronic Enh) are the *cis*-regulatory element (CRE) candidates. **B, C**. qRT-PCR analysis of mRNA levels of UGT1A isoforms (UGT1A3, 1A4, 1A6, 1A7 and 1A9) in rifampicin (Rif)-treated ΔPBREM 1, 2 and ΔPBREM-like 1, 2 (B) or Rif-treated ΔIntronic enh. 1-3 (C). The expression levels are normalized to *GAPDH* mRNA. All experiments were performed in triplicate or quadruplicate for each condition and repeated at least three times with similar results. Representative data were shown. The error bars indicate standard deviation, and unpaired Welch's *t*-test was used to calculate the *P* value. $*P < 0.05$, $**P < 0.01$, and $***P < 0.001$.

Fig. S4

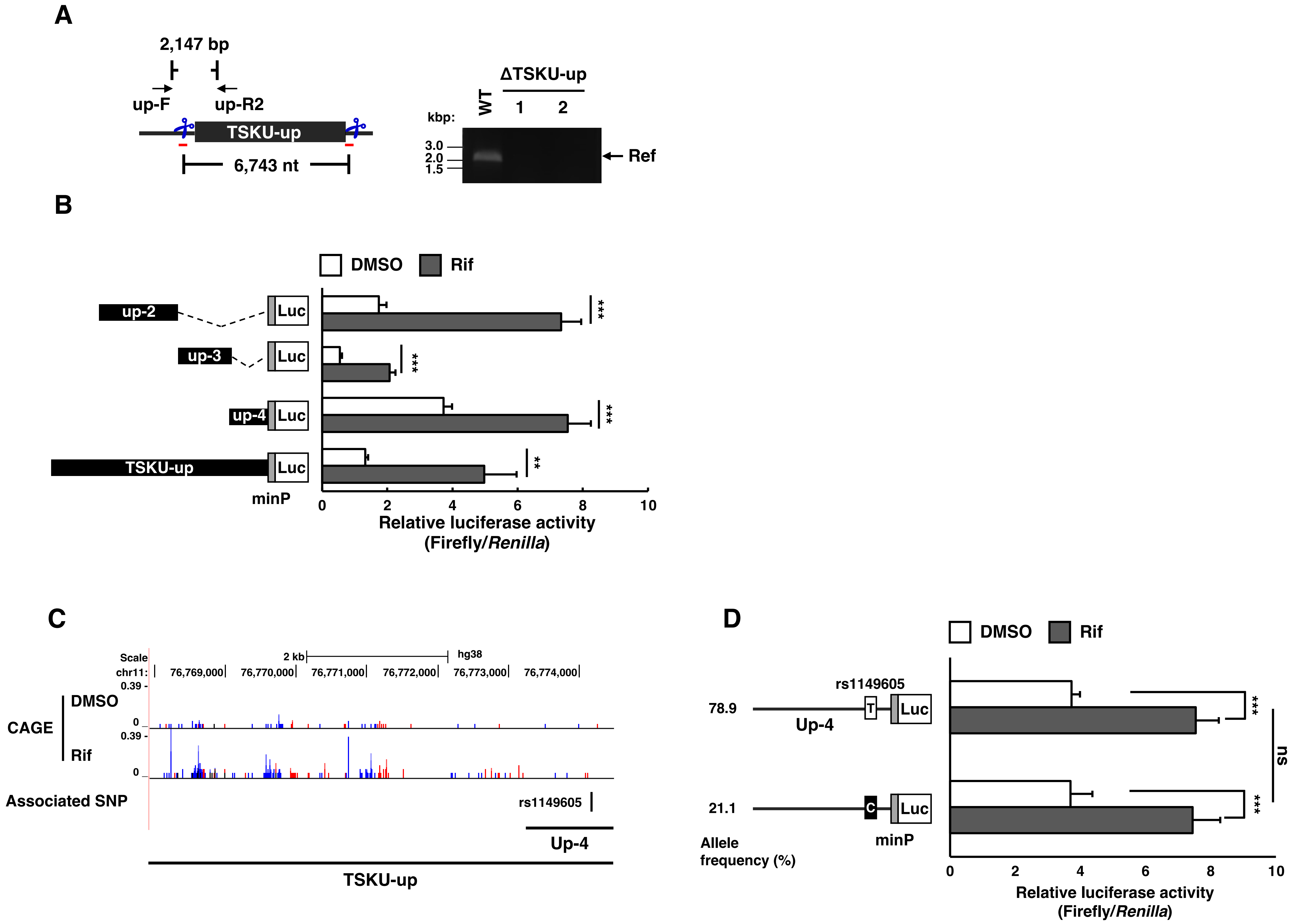


**Fig. S4**
**A** Schematic view of CRISPR/Cas9-based deletion of TSKU-up in ShP51 cells with PCR primers used for amplification of genomic DNA (Left). Gel images show the PCR products from genomic DNA extracted from the deletion mutants (ΔTSKU-up 1, 2) (Right). **B** Luciferase activity of each of Up-2, -3, -4 and TSKU-up with the minimal promoter (minP) in rifampicin (Rif)-treated ShP51 cells. **C** A view of the TSKU-up region (chr11:76,767,944-76,774,353) in UCSC Genome Browser . up-4 contains GWAS SNP, rs1149605 associated with vitamin D levels. **D** Effect of rs1149605 allele on luciferase activity of up-4 with minP in Rif-treated ShP51 cells. In luciferase reporter assays, Firefly luciferase activity was normalized to *Renilla* luciferase activity. Results are expressed as fold change compared with empty vector control. All experiments were performed in triplicate for each condition and repeated at least three times with similar results. Representative data were shown. The error bars indicate standard deviation, and unpaired Welch's *t*-test was used to calculate the *P* value. ****P* < 0.001. ns, not significant

Fig. S5

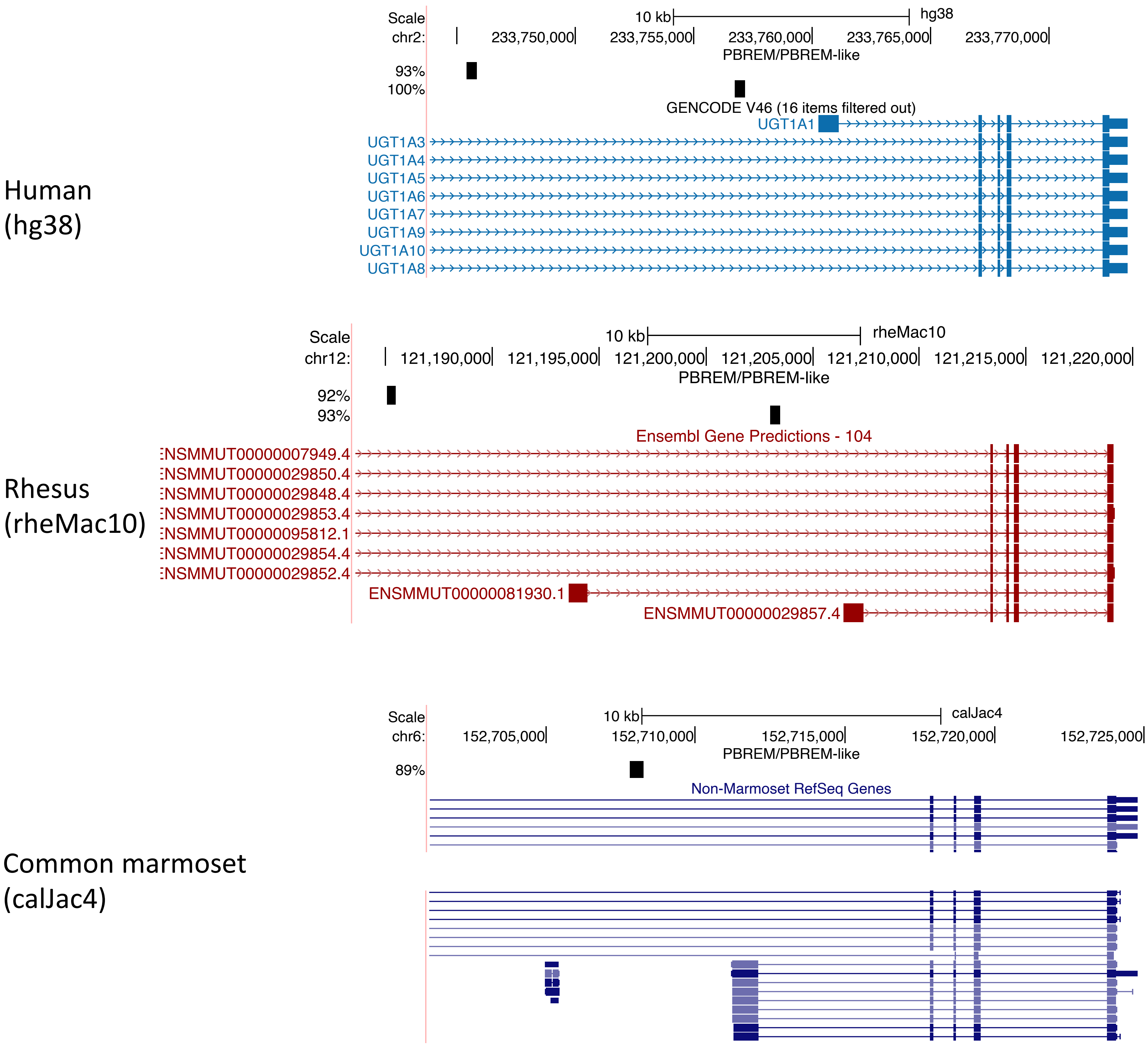


**Fig. S5**
**Homologous sequences of PBREM in primates.**
The DNA sequence corresponding to the PBREM (or DPE16, located at chr2:233756748-233757190 on hg38) was aligned with three primate species with BLAT. Two regions were identified in human and rhesus macaque (old world monkey), while a single region found in common marmoset (new world monkey). No corresponding sequence were found in mouse.
